# Spatial Ecology of Territorial Populations

**DOI:** 10.1101/694257

**Authors:** Benjamin G. Weiner, Anna Posfai, Ned S. Wingreen

## Abstract

Many ecosystems, from vegetation to biofilms, are composed of territorial populations that compete for both nutrients and physical space. What are the implications of such spatial organization for biodiversity? To address this question, we developed and analyzed a model of territorial resource competition. In the model, all species obey trade-offs inspired by biophysical constraints on metabolism; the species occupy non-overlapping territories while nutrients diffuse in space. We find that the nutrient diffusion time is an important control parameter for both biodiversity and the timescale of population dynamics. Interestingly, fast nutrient diffusion allows the populations of some species to fluctuate to zero, leading to extinctions. Moreover, territorial competition spontaneously gives rise to both multistability and the Allee effect (in which a minimum population is required for survival), so that small perturbations can have major ecological effects. While the assumption of trade-offs allows for the coexistence of more species than the number of nutrients – thus violating the principle of competitive exclusion – overall biodiversity is curbed by the domination of “oligotroph” species. Importantly, in contrast to well-mixed models, spatial structure renders diversity robust to inequalities in metabolic trade-offs. Our results suggest that territorial ecosystems can display high biodiversity and rich dynamics simply due to competition for resources in a spatial community.

---

Living things exist not in isolation but in communities, many of which are strikingly diverse. Tropical rainforests can have more than 300 tree species in a single hectare [1], and it has been estimated that one gram of soil contains 2,000-30,000+ distinct microbial genomes [2, 3]. Understanding the relationship between biodiversity and the environment remains a major challenge, particularly in light of the competitive exclusion principle: in simple models of resource competition, no more species can coexist indefinitely than the number of limiting resources [4, 5]. In modern niche theory, competitive exclusion is circumvented by mechanisms which reduce niche overlaps and/or intrinsic fitness differences [6, 7], suggesting that trade-offs may play an important role in the maintenance of biodiversity. Intriguingly, diversity beyond the competitive-exclusion limit was recently demonstrated in a resource-competition model with a well-mixed environment and exact metabolic trade-offs [8]. However, many ecosystems are spatially structured, and metabolic trade-offs are unlikely to be exact. While some spatial structure is externally imposed, it also arises from the capacity of organisms to shape their environment. How does self-generated spatial structure, along with realistic metabolic constraints, impact diversity?

Various studies have clarified how intrinsic environmental heterogeneity (e.g. an external resource gradient) fosters biodiversity by creating spatial niches [9–13]. Others have demonstrated that migration between low-diversity local environments can lead to “metacommunities” with high global diversity [14–19]. But how is diversity impacted by *local* spatial structure? Recent models suggests that spatial environments without intrinsic heterogeneity can support higher diversity than the well-mixed case [20–25], although the effect depends on the interactions and details of spatial structure [26, 27]. In these models, competition follows phenomenological interaction rules. In some cases, tradeoffs have been invoked to limit fitness differences [21] and penalize niche overlap [25], but did not otherwise structure the spatial interactions. All these models allow coexistence when the combination of spatial segregation and local interactions weakens interspecific competition relative to intraspecifc competition. However, it remains unclear how such interactions relate to concrete biophysical processes.

Here, we study biodiversity in a model where species interact through spatial resource competition. We specifically consider surface-associated populations which exclude each other as they compete for territory. This is an appropriate description for biofilms, vegetation, and marine ecosystems like mussels [28] or coral [29], in contrast with models that represent populations as overlapping densities and better describe motile or planktonic populations [9, 30]. The well-mixed environment is an explicit limit of our model, so we are able to isolate the unique effects of spatial structure.

We find that, contrary to expectations, introducing population territories into a model with metabolic trade-offs *reduces* biodiversity relative to the well-mixed case. Extinctions occur over a new timescale inversely related to the nutrient mixing time. Spatial structure also leads to the emergence of multiple steady states and the Allee effect, so that small perturbations may have drastic consequences. Finally, we find that overall biodiversity is curbed by the domination of “oligotroph” species but is robust to inequalities in metabolic tradeoffs.

## RESULTS

### Model

We developed a model of territorial populations competing for diffusing resources to clarify the relationship between spatial structure, metabolic trade-offs, and biodiversity. The model is spatially explicit and relates the mechanistic dynamics of competition to parameters with clear biological meaning. Crucially, competing populations are not interpenetrating, so populations are competing for both nutrients and territory.

Specifically, we consider *m* species competing for *p* nutrients in a one-dimensional space of size *L* with periodic boundary conditions (a ring). The rate of supply of nutrients is specified by the supply vector 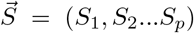 such that Σ_*i*_*S_i_* = *S*, where *S* is the total nutrient supply rate in units of concentration/time. The nutrient supply is spatially uniform, so there is no external environmental heterogeneity. Each species *σ* ∈ [1…*m*] is defined by its metabolic strategy 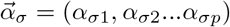, which specifies the proportion of its metabolic resources (e.g. enzymes) it allocates to the consumption of each nutrient. Metabolic trade-offs are implemented via a constraint on the enzyme budget, namely Σ_*i*_*α*_*σi*_ = *E* for all species (except where noted). Metabolic strategies and the supply can be represented as points on a simplex of dimension *p* − 1 (Fig. 1*A* and Fig. 2*A* inset, for example). Each species occupies a segment of the ring corresponding to its population *n_σ_*, so that *n_σ_* is a length and *σ* = 1…*m* specifies a spatial ordering. For example, the population with strategy 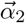 occupies the segment of the ring between populations with strategies 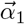 and 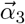. Populations never overlap, so the total population satisfies Σ_*σ*_ *n_σ_* = *L*. Figures 1*B* and *C* show an example of the time evolution of one such spatial community consisting of 11 species competing for three nutrients.

**Figure 1.**
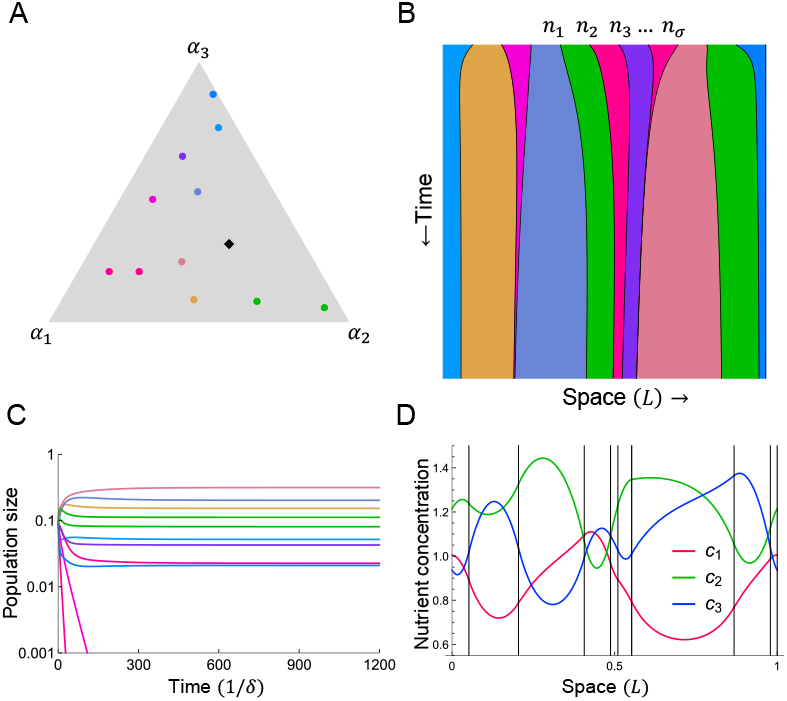
A model with spatial structure and metabolic trade-offs supports more species than expected from the principle of competitive exclusion. Example with three nutrients and eleven species starting with equal populations. (*A*) Each species uptakes nutrients according to its enzymeallocation strategy (*α*_*σ*1_, *α*_*σ*2_, *α*_*σ*3_). Because strategies satisfy the budget constraint Σ_*i*_ *α_σi_* = *E*, each can be represented as a point on a triangle in strategy space. The nutrient supply 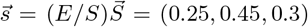 is represented as a black diamond. Colors correspond to strategies and are consistent throughout the figure. (*B*) Each species occupies a fraction of a one-dimensional space (a ring) and has a corresponding time-dependent population size *n_σ_* (*t*). Here, the nutrient diffusion time *τ_D_* is 400. (*C*) Population dynamics from *A*. Nine species coexist on three nutrients. (*D*) Concentrations of the three nutrients at steady state (vertical black lines denote boundaries between populations).

While the supply of nutrients is spatially uniform, the local rate of nutrient consumption depends on the metabolic strategy of the local species, and nutrients diffuse in space. We study the regime where population growth is nutrient-limited, so the rate of uptake of each nutrient is linear in its concentration. Thus, within each region occupied by a single species *σ*, the nutrient concentrations *c_σi_* obey

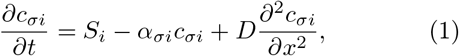

where *D* is the diffusion coefficient for all nutrients. As nutrient processing is generally much faster than growth, we assume a separation of timescales, such that nutrient concentrations equilibrate before populations change. Then 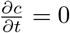, and

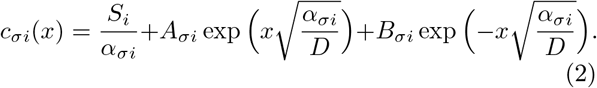

The constants of integration *A_σi_* and *B_σi_* are fixed by the physical requirement that *c_i_*(*x*) be continuous and differentiable at the population boundaries. Figure 1*D* shows the concentrations of the three nutrients after the populations shown in Fig. 1*B* and *C* have reached steady state. The competitors transform the uniform nutrient supply into a complex spatial environment by depleting their preferred nutrients while allowing other nutrients diffuse to their neighbors.

The populations change in time according to

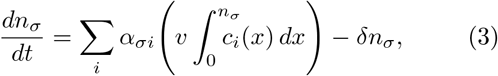

where *δ* is the death rate and the integral is taken over the territory occupied by species *σ. v* is a length that converts nutrients to territory growth. The total population remains fixed at *L*, corresponding to competition for a share of a fixed total territory. This implies 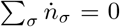, which requires *δ* = *vS*, i.e. the death rate matches the nutrient value of the total supply rate. We choose units of time and concentration such that *v* = 1 and *S* =1, without loss of generality. In the example shown in Fig. 1*B* and *C*, nine species coexist, far exceeding the three-species limit set by competitive exclusion.

The spatial nutrient environment influences the population dynamics via the dimensionless diffusion time *τ_D_* ≡ *L*^2^*E/D*, which is the time for nutrients to diffuse a distance *L* relative to the uptake time. Competitors interact only through the nutrient environment, so when nutrients diffuse instantaneously (*τ_D_* = 0), the spatial dynamics reduce to the well-mixed dynamics. (See *SI Appendix* for the *τ_D_* → 0 expansion.)

### Biodiversity

How does territorial spatial structure influence biodiversity? As an illustrative example, we consider ten species competing for two resources. The simplex in the inset of Fig. 2*A* shows how each of the strategies (colored dots) and the nutrient supply (diamond) divide between the two nutrients. In Fig. 2*A*, the nutrients are well-mixed (*τ_D_* = 0), and all ten species coexist at steady state. The steady state of the spatial case shown in Fig. 2*B* still exceeds competitive exclusion, with three species coexisting on two resources, but much of the biodiversity is lost. This behavior is striking, as it contrasts with many competition models where spatial structure *increases* diversity relative to the well-mixed case [20–26]. In those models, diversity increases because spatial segregation, combined with local interactions, weakens interspecific competition. Here, however, the resource environment is uniformly coupled via diffusion, so competition remains strong. Strategies that are poorly matched to the nutrient supply allow unused nutrients to diffuse away to competitors; such populations shrink until the nutrient fluxes are balanced or the species goes extinct. This contrasts with the well-mixed case, where at steady state all the nutrient concentrations are equal so every strategy can coexist [8]. Thus territorial spatial structure heightens competitive differences between strategies, even when all obey the same trade-offs.

**Figure 2.**
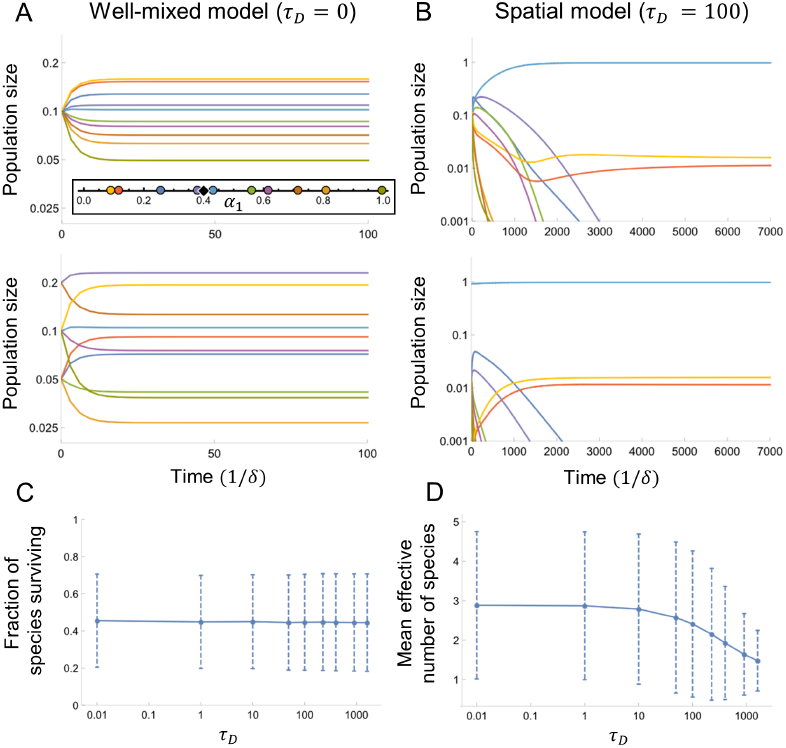
Spatial structure reduces diversity compared to the well-mixed limit of instantaneous nutrient diffusion. (*A*) Top: A well-mixed population of ten species with equal initial populations competing for two nutrients. All ten coexist at steady state. Bottom: Same as Top, but with different initial populations. The community reaches a new steady state. (Inset) Strategies 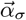 and resource supply 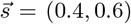. (*B*) Top: Same species and nutrient supply as *A*, but in a spatial environment with nonzero nutrient diffusion time. Only three species survive. Bottom: Same as Top, but with different initial populations. The community reaches the same steady state. (*C*) Fraction of initial species coexisting at steady state with 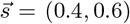; a population is considered extinct if *n_σ_/L* < 10^−6^ (mean ± SD for 400 random sets of ten strategies). The well-mixed model has survival fraction 0.99±0.08. (D) Effective number of species *M* at steady state (mean ± SD for same strategies as *C*).

How representative is the behavior seen in Fig. 2*A* and *B*? In Fig. 2*C* and *D* we show results for many randomly generated territorial communities, confirming that the loss of biodiversity is a generic feature of spatial structure, and that the nutrient diffusion time *τ_D_* acts as a control parameter for biodiversity. In the well-mixed model, all ten species typically coexist. (See [8] for a discussion of the “convex hull condition” for coexistence.) Figure 2*C* shows the mean fraction of species coexisting at steady state for nonzero *τ_D_*. Spatial communities still violate competitive exclusion, but a large fraction of species go extinct. Even among those that survive, spatial structure reduces biodiversity by rendering abundances highly unequal. Using the same data as *C*, Fig. 2*D* quantifies this via the effective number of species *M* = exp{(*H*)}, where *H* = – Σ_*σ*_ *p_σ_* log*p_σ_* is the Shannon entropy and *p_σ_* = *n_σ_*/*L*. (Intuitively, *M* is the number of equal populations yielding *H*. See *SI Appendix* for full rank abundance curves.) The average community loses ≈ 1/3 of its steady-state diversity as *τ_D_* grows from 0.01 – 1600. In well-mixed communities, all ten species are typically present in comparable proportions, but in the spatial model, one species dominates. Once this population is large compared to 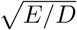, increasing *τ_D_* adds to the “bulk” population in its interior, decreasing overall diversity (see *SI Appendix* for details). However, aggregate measures of diversity in 2*C* and *D* belie a wide distribution of outcomes. For example, only three species survive in Fig. 2*B*, whereas nine coexist in Fig. 1. Why are some steady-state communities so much more diverse than others?

In order to identify which features of the initial set of species determine steady-state diversity, we generated many random communities with species drawn uniformly from strategy space. Figures 3*A* and *C* show the distributions of the steady-state diversity *M* as a function of *s*_1_, the supply of Nutrient 1. The number of nutrients does not explain the difference in outcomes. However, diverse steady states proliferate as the supply becomes more balanced between nutrients. What distinguishes high diversity outcomes? Figures 3*B* and *D* show every strategy present in every community with high diversity. Diverse communities have one thing in common: they lack species in the region of strategy space where 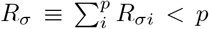. Here, *R_σi_* ≡ *S_i_*/*α_σi_* is the uniform concentration of Nutrient *i* an isolated population with uptake *α_σi_* would produce given a supply rate *S_i_*. Thus *R_σ_* is the total nutrient concentration maintained by and sustaining an isolated species *σ* at steady state. For comparison, *R_σ_* diverges for specialists (*α_σi_* = 0), while a perfect generalist (*α_σi_* = 1 /*p*) has *R_σ_* = *p*, as does a strategy that perfectly matches the supply (*α_σi_* = *S_i_*). Strategies satisfying *R_σ_* < *p* survive on even *lower* total nutrient concentrations, so we christen them “oligotrophs.” Their ability to create and survive on the minimum total nutrient concentration allows them to drive competitors extinct, thus reducing diversity. This recalls Tilman’s famous result that the species with the lowest equilibrium concentration of its limiting resource (the lowest *R*\*) can displace all others competing for that resource [15]. However, the *R*\* rule is due to a species’ innate superiority in consuming a single resource, whereas the oligotroph condition arises in a competition for multiple resources between intrinsically equal species.

**Figure 3.**
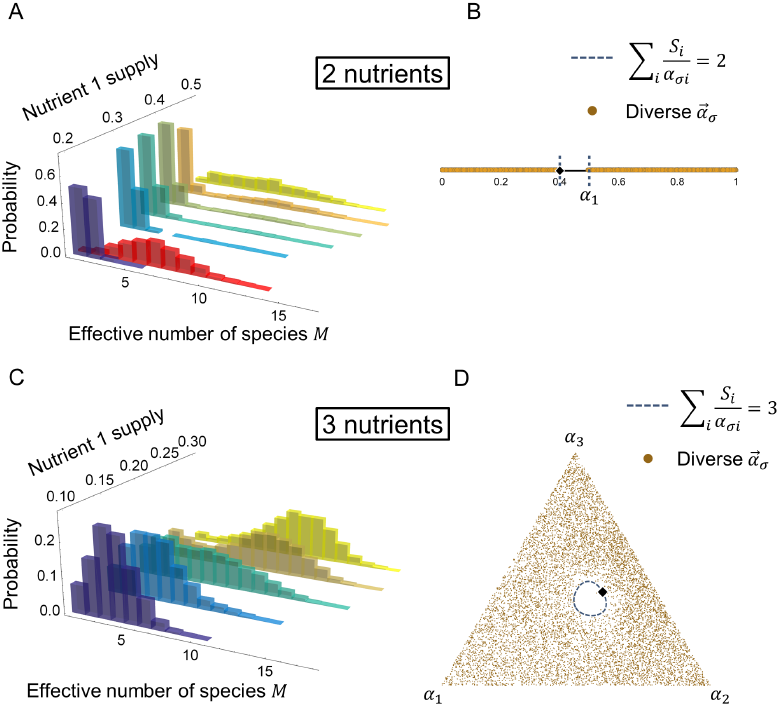
Steady-state diversity is governed by a simple condition: diversity crashes if there is an “oligotroph” whose strategy satisfies *R_σ_* < *p*. (*A*) Probability of effective number of species *M* at steady state. For each nutrient supply, we simulated 2000 sets of 20 strategies. Strategies were chosen uniformly at random except the case shown in red 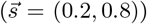, where oligotrophs were excluded. *τ_D_* = 10 here and below. (*B*) For 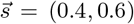 (orange in *A*), we plot all strategies that appear in the most diverse 10% of simulations (90th percentile and above of *M*). No strategies appear in the oligotroph region, demarcated by the blue dashed lines. (*C*) Same as *A* but for three nutrients. (*D*) Strategies that appear in the most diverse 10% of simulations for 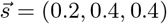 (teal in *C*). The oligotroph region is nearly empty.

To test whether it is simply the presence/absence of oligotrophs that controls overall biodiversity, we generated random communities in an environment with an asymmetric nutrient supply (*s*_1_ = 0.2), but excluded oligotrophs. The resulting steady-state communities are much more diverse (Fig. 3*A*, red) than the case where oligotrophs are allowed (Fig. 3*A*, purple). Hence an asymmetric nutrient supply reduces diversity by increasing the probability that an oligotroph will be present. The oligotroph condition captures the intuition of the *R*\* rule – species with low resource requirements dominate – but does not require biological superiority or preclude coexistence beyond competitive exclusion.

### Alternative Steady States and Slow Dynamics

How does the outcome of spatial competition depend on initial conditions? Consider the well-mixed system in Fig. 2*A*. All that differs between the top and bottom subplots are the initial populations, but the same set of species has two very different steady states; not even the hierarchy of populations is preserved. In fact, there is an *m* − *p* dimensional degenerate manifold of fixed points corresponding to the communities that construct the same steady-state nutrient environment 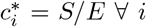. The final population may lie anywhere on this manifold. By contrast, in the spatial ecosystem of Fig. 2*B*, both sets of initial populations converge to the same unique steady state.

The relationship between the steady states in the well-mixed and spatial regimes can be visualized in a simple example. Figure 4*A* shows the phase behavior of three species competing for two resources. Here, *m − p* = 1, so the well-mixed case (left) has a onedimensional degeneracy of steady states. In the spatial community (middle/right), the degenerate manifold collapses to a single fixed point. (Here the fixed point is unique, but this is not always the case; see Fig. 5.) This discontinuous change in the steady states is reflected in Fig. 2*C* and *D*, where the diversity for any *τ_D_* ≠ 0 is substantially lower than for the well-mixed limit *τ_D_* = 0. Figure 4*A* also clarifies another striking difference between Fig. 2*A*, in which the well-mixed community approaches steady state at approximately the individual death rate *δ*, and Fig. 2*B*, in which the spatial community approaches steady state orders of magnitude more slowly. This emergent slow timescale and the breaking of degeneracy are intimately related: for any nonzero diffusion time *τ_D_*, the degenerate manifold becomes a corresponding slow manifold, which the population rapidly reaches and then crawls to a fixed point. Linear stability analysis around this fixed point reveals a relaxation time *t*_slow_ ~ 1/*τ_D_*, which diverges as *τ_D_* → 0. (See *SI Appendix* for details.)

**Figure 4.**
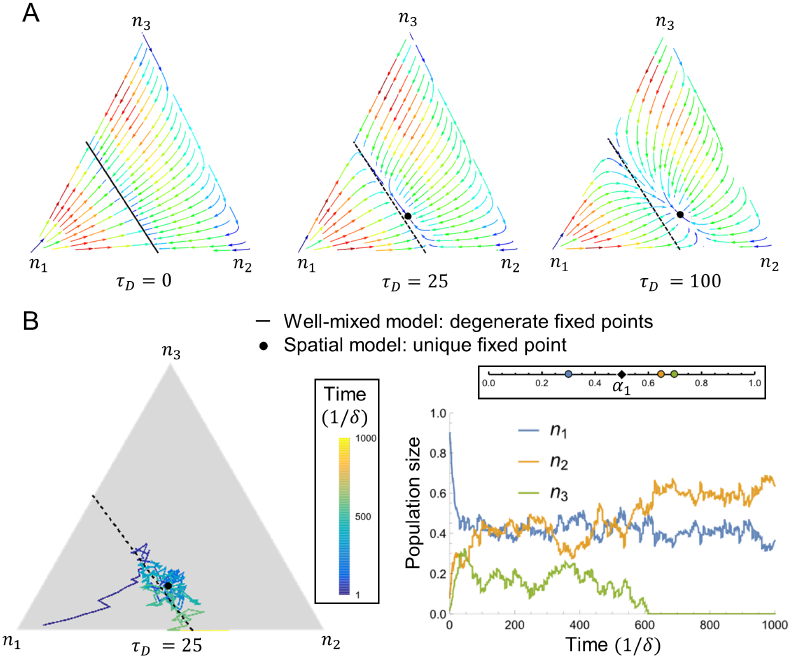
Spatial structure replaces the steady-state degeneracy of the well-mixed case with slow modes in population space. (*A*) Trajectories in population space for a three-way competition at different values of *τ_D_* (Inset: strategies and supply). The direction and color of the arrows show the direction and magnitude of *dn_σ_/dt*, respectively. (B) Same as *A*, but with stochastic dynamics due to random births and deaths; see *SI Appendix* for details. (Inset: Trajectory color as a function of time.) *Left*: Species 3 drifts to extinction. *Right: n_σ_*(*t*) for the population trajectory at *Left*.

What are the ecological implications of this slow relaxation to steady state? In general, diverse communities with *m − p* ≫ 1 could have tens or hundreds of slow modes for population changes. These modes shape the response to perturbations: a microbial community might recover from one antibiotic very rapidly and another very slowly, depending on the shift in population space. Even without an intervention, real populations will have stochastic fluctuations around the steady state. Figure 4*B* shows trajectories through population space for the same species and nutrient supply as in Fig. 4*A*, but with demographic noise due to stochastic births and deaths. Ecological drift is confined to the slow manifold, and fluctuations primarily excite the population’s “soft mode” of the balance between Species 2 and Species 3. These two have similar strategies, and either can drift to extinction, whereas Species 1 always survives. In the absence of noise, increasing *τ_D_* decreases fixed-point diversity (Fig. 2*D*). With noise, however, steady states in the well-mixed limit are unstable to fluctuations along the degenerate manifold. Increasing the “restoring force” (~ *τ_D_*) can prevent species from fluctuating to extinction, and so spatial structure can stabilize diversity.

Although the well-mixed case has degenerate steady states, the steady-state nutrient environment is unique, and small initial population differences lead to small differences in the steady state (see Fig. 4*A*). By contrast, spatial communities can have multiple steady-state nutrient environments, and similar populations may end in very different steady states. For example, in a competition of two species for two resources, Fig. 5*A* and *B* show the steady states as a function of *α*_21_, with *α*_11_ held fixed. (*α*_*σ*1_ is the enzyme allocation of Species *σ* to Nutrient 1. Due to trade-offs, this also fixes *α*_*σ*2_.) In Fig. 5*A*, there are two alternative steady states with both species coexisting. The unstable fixed point separates the relatively equal community of the lower branch from the upper branch where Species 1 dominates. This bistability leads to discontinuous transitions, where small changes (the populations crossing the separatrix, or the strategy exiting the bistable phase) can have dramatic consequences. Figure 5*B* shows another region of strategy space where the outcomes are bistable, but now the alternatives are coexistence and exclusion. Above the unstable fixed point in Fig. 5*B*, Species 1 drives Species 2 to extinction. Otherwise, they coexist with Species 2 having the larger population. This is an example of the Allee effect: Species 2 can only survive if its population exceeds a threshold. (See *SI Appendix* for a phase diagram of the full strategy space.)

**Figure 5.**
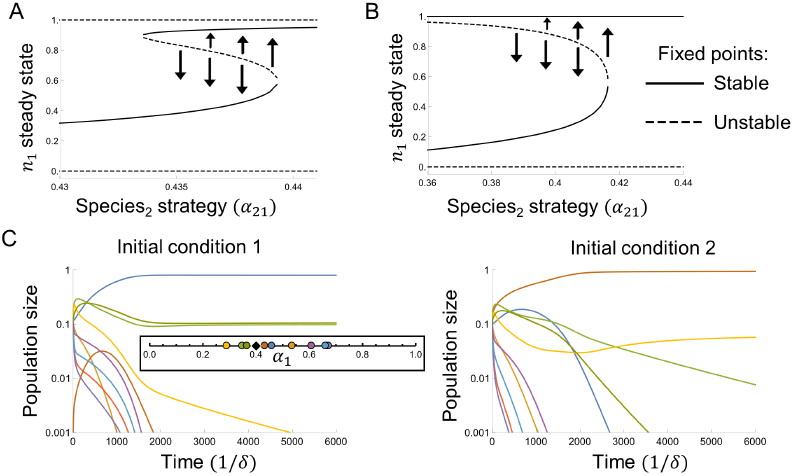
(*A, B*) Two species competing for two nutrients, with supply 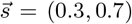 and *τ_D_* = 400. Arrows indicate flow away from unstable fixed points. (*A*) Bistability for *α*_11_ = 0.29. (*B*) Allee effect for *α*_11_ = 0.31. Species 2 goes extinct if its initial population is too low. (*C*) The Allee effect in a competition with ten species and two nutrients, with 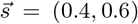 and *τ_D_* = 100. The blue and brown species displace each other, depending on initial conditions.

The Allee effect persists in more complex communities. Figure 5*C* shows a ten-species competition where the brown and blue species can displace each other depending on the initial conditions, modifying the eight other species’ fates in the process. Thus multistability and the Allee effect emerge naturally in our territorial model, even though the species interact exclusively through competition for resources.

### Unequal Enzyme Budgets

Metabolic trade-offs are plausible because all microbes face the same biophysical constraints on metabolism and protein production, but trade-offs are unlikely to be exact in real ecosystems. How does this impact biodiversity in our model? Figure 6*A* shows results for ten species with exact trade-offs (Σ*_i_ α_σi_* = *E* for all species) competing for two resources. The well-mixed community is very diverse, while the spatial community is not. In Fig. 6*B*, each species allocates the same fraction of its enzyme budget to each nutrient as in 6A, but each with its own total enzyme budget *E_σ_*. Diversity collapses in the well-mixed system, but the spatial community actually becomes more diverse. Figures 6*C* and *D* show that this behavior is typical via comparison of the steady-state diversity of communities where each species’ enzyme budget is drawn from a Normal Distribution with mean 1 and standard deviation *δE*. Well-mixed communities (*C*) are very diverse if trade-offs are exact (*δE* = 0), but any disparity in the enzyme budgets causes diversity to collapse; only one or two species survives at steady state. The spatial communities (*D*) are less diverse for *δE* = 0, but their diversity 〈*M*〉 actually increases with *δE*. This is due to an asymmetric effect: oligotrophs lose their dominance with a very small decrease in *E_σ_*, but other species require a large increase in *E_σ_* to dominate (see *SI Appendix* for details). As a result, spatial communities with imperfect trade-offs can display biodiversity well beyond the competitive-exclusion limit.

**Figure 6.**
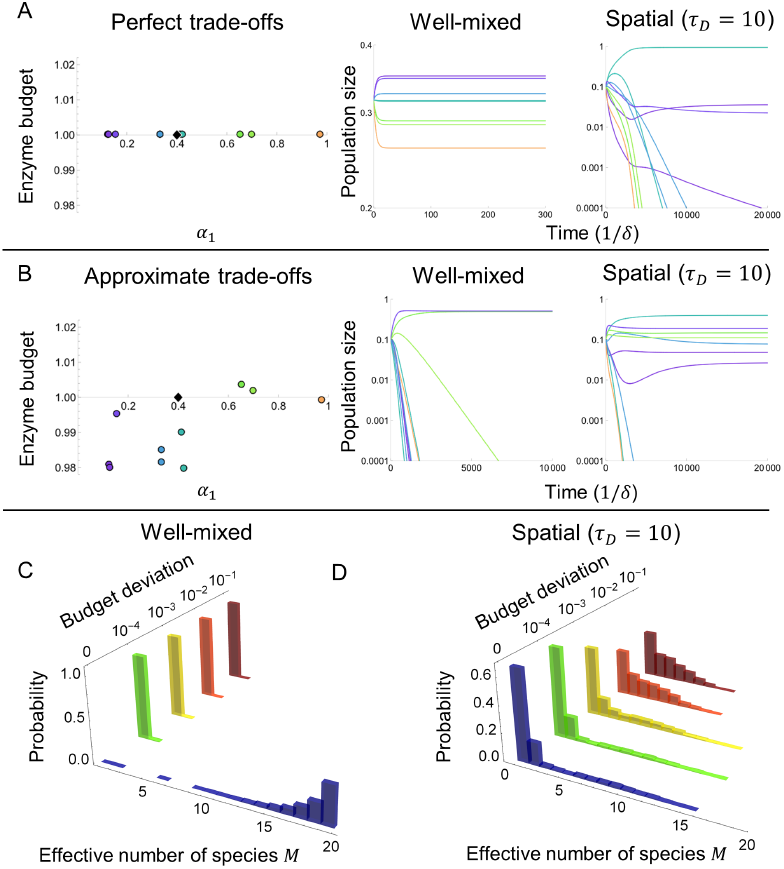
Territorial spatial structure renders diversity robust to variation in enzyme budgets. (*A*) In a community with equal enzyme budgets and 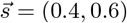 (*Left*), ten species coexist in the well-mixed model (*Center*), whereas only three coexist in the spatial model (*Right*). (*B*) In a community with the same strategies as *A* but unequal enzyme budgets (*Left*), two species coexist in the well-mixed model (*Center*), whereas seven coexist in the spatial model (*Right*). (*C*) Probability of effective number of species *M* at steady state for random enzyme budgets, in the well-mixed model with 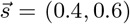. For 2000 sets of strategies, each of 20 initial species’ enzyme budgets *E_σ_* was drawn from 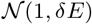. (*D*) Same as *C* but for spatial model.

## DISCUSSION

We analyzed a model of spatial resource competition among territorial surface communities such as biofilms, vegetation, or coral. Each species has a concrete metabolic strategy subject to biophysical tradeoffs. The nutrient environment has no intrinsic heterogeneity but is globally coupled via diffusion, so competitors shape it via consumption. We found that the resulting spatial structure restricts biodiversity, in stark contrast to previous models where spatial segregation increases diversity by weakening competition. In the simplest of these cases, different resources are partitioned into different regions, providing spatial niches [9–13]. Alternatively, competitors may self-organize into patches linked by migration [14–16, 18, 19] or into aggregates with local interactions [20–26]. External resource gradients can also increase diversity, because diffusion of dense motile populations prevents any species from monopolizing resource-rich regions [9, 10]. In our model, the situation is very different. Because the external nutrient supply is uniform and the entire space is linked via diffusion, no spatial niches emerge; competitors have nowhere to hide. This is reminiscent of ecological reaction-diffusion models without external sources, where global coupling reduces diversity [26] and nonuniform steady states only become possible for unequal diffusion coefficients or complex geometries [27]. Our communities are fundamentally different, however, as they occupy exclusive territories and exceed the competitive-exclusion limit in spite of a simple geometry and uniform diffusion coefficients.

What controls diversity in our model? The degree of nutrient mixing *τ_D_* controls the evenness of abundances by setting the population of the dominant species, while the presence of oligotrophs distinguishes steady states of high coexistence from those with many extinctions. Oligotrophs drive competitors extinct because they have the lowest total nutrient requirements, in rough analogy with the lowest *R\** rule for well-mixed systems [15]. However, oligotrophs obey the same trade-offs as every other species, and their dominance arises from the relationship between their strategies and the nutrient supply rather than any innate superiority. The composition of the nutrient supply sets the strategy range of oligotrophs, so it is effectively another control parameter for diversity. In spite of highly nonlinear dynamics and many parameters, the oligotroph condition provides a simple criterion for diversity.

Spatial structure also provides a novel mechanism for discontinuous transitions between alternative steady states. Such sudden shifts, or “catastrophes”, attract significant attention due to their implications for ecosystem resilience [31]. The Allee effect occurs in a large variety of ecosystems [32], and is particularly relevant to the conservation of rare species. It is usually understood as the result of transparently cooperative processes, such as production of a public good [32], and modeled via an explicit cooperative term. In resource-competition models, multistability has been observed when species consume nutrients one at a time [33] or with unequal stoichiometries [34]. Here, both the Allee effect and multistability emerge naturally from the ability of a population to render its resource environment more favorable to itself. Interestingly, the Allee-effect species are oligotrophs, underscoring the special ability these strategies have to impact their ecosystems.

It has been observed that spatial structure increases the time to reach equilibrium [35]. Here we showed precisely how a new dynamical timescale emerges from spatial structure. We found that the slow dynamics are confined to a manifold in population space. These slow modes of the population are subject to large fluctuations due to noise (e.g. demographics). Slow relaxation also means that for a rapidly changing nutrient supply, the population might never reach steady state, potentially saving some species from extinction.

Finally, we find that spatial structure allows diversity to persist with imprecise metabolic trade-offs. In the well-mixed system without noise, any deviation from exactly equal enzyme budgets leads to ecosystem collapse [8]. Spatial communities, however, remain diverse with only approximate trade-offs. In fact, variation in enzyme budgets actually increases mean diversity by impairing oligotrophs. The persistence of diversity beyond competitive exclusion with inexact trade-offs makes it more credible that trade-offs play a role in maintaining the surprising diversity of real ecosystems.

Our results suggest several future research directions. A two-dimensional extension of the model exhibits the same loss of biodiversity due to oligotrophs and uneven abundances (see *SI Appendix*), and it will be interesting to explore 2D pattern formation in more depth. One might also consider resources that diffuse at different rates. This can lead to nonuniform steady states in reaction-diffusion systems [27]. Finally, in microbial communities, gene regulation and evolution are often relevant on ecological timescales, so it would be natural to allow species to modify their strategies.

In summary, we find that spatial structure engenders more realistic communities: it curtails the unlimited diversity of the well-mixed model, but allows for coexistence beyond the competitive exclusion principle even in the absence of exact metabolic trade-offs. Our results demonstrate that mechanistic interactions, arising from biophysical constraints such as space and metabolism, can allow even simple models to capture some of the rich behaviors of real ecosystems.

## METHODS

The population ODEs (Eq. 3) were solved numerically using Mathematica’s “NDSolve”. The *c_σ,i_* depend on *n_σ_* through the coefficients {*A_σi_,B_σi_*}, which are fixed by requiring that *c_i_*(*x*) be continuous and differentiable at the population boundaries. The nutrient equations are simpler under the change of variables 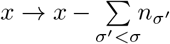. Then *c_σ,i_*(*x*) runs from 0 to *n_σ_*, yielding the system

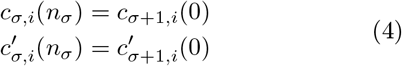

which was solved using Mathematica’s “LinearSolve” with periodic boundary conditions (*c_mi_*(*n_m_*) = *c*_1*i*_(0), corresponding to a ring).

Details on the well-mixed model, stochastic dynamics, and figure parameters can be found in *SI Appendix*. Code will be available on GitHub.

## ACKNOWLEDGEMENTS

We thank S. Levin for valuable conversations and suggestions. This work was supported in part by the National Science Foundation, through the Center for the Physics of Biological Function (PHY-1734030) (B.G.W.), the National Institutes of Health (www.nih.gov), Grant R01 GM082938 (N.S.W.), and the Princeton University Diekman Genomics Fund (A.P.).

## References

[1] A. H. Gentry, Proc Natl Acad Sci USA 85, 156 (1988).

[2] R. Daniel, Nature Reviews Microbiology 3, 470–478 (2005).

[3] T. P. Curtis, W. T. Sloan, and J. W. Scannell, Proc Natl Acad Sci USA 99, 10494–10499 (2002).

[4] R. Macarthur and R. Levins, Proc Natl Acad Sci USA 51, 1207 (1964).

[5] S. A. Levin, The Am. Nat. 104, 413 (1970).

[6] P. Chesson, Annual Review of Ecology and Systematics 31, 343–366 (2000).

[7] J. M. Chase and M. A. Leibold, Ecological Niches (University of Chicago Press, 2003).

[8] A. Posfai, T. Taillefumier, and N. S. Wingreen, Physical Review Letters 118 (2017).

[9] S.-B. Hsu and P. Waltman, SIAM Journal on Applied Mathematics 53, 1026 (1993).

[10] J. Huisman, P. Oostveen, and F. J. Weissing, The Am. Nat. 154, 46 (1999).

[11] M. Doebeli and U. Dieckmann, Nature 421, 259 (2003).

[12] D. Tilman, Proc Natl Acad Sci USA 101, 10854 (2004).

[13] A. B. Ryabov and B. Blasius, Ecology Letters 14, 220 (2011).

[14] S. A. Levin, Annual Review of Ecology, Evolution, and Systematics 7, 287 (1976).

[15] D. Tilman, Ecology 75, 2 (1994).

[16] M. Loreau, N. Mouquet, and R. D. Holt, Ecology Letters 6, 673 (2003).

[17] J. M. Kneitel and J. M. Chase, Ecology Letters 1, 69 (2004).

[18] D. Gravel, C. D. Canham, M. Beaudet, and C. Messier, Ecology Letters 9, 399 (2006).

[19] L. G. Shoemaker and B. A. Melbourne, Ecology 97, 2436 (2016).

[20] D. J. Murrell and R. Law, Ecology Letters 6, 48 (2003).

[21] C. Dislich, K. Johst, and A. Huth, Eco. Modelling 221, 2227 (2010).

[22] N. Mitarai, J. Mathiesen, and K. Sneppen, Physical Review E 86 (2012).

[23] J. Vandermeer and S. Yitbarek, Journal of Theoretical Biology 300, 48 (2012).

[24] J. Velázquez, J. P. Garrahan, and M. P. Eichhorn, PLoS ONE 9 (2014).

[25] P. V. Martín, J. Hidalgo, R. Rubio de Casas, and M. A. Muñoz, PLOS Computational Biology 12 (2016).

[26] R. Durrett and S. A. Levin, Theoretical Population Biology 46, 363 (1994).

[27] S. A. Levin, in Pattern Formation by Dynamic Systems and Pattern Recognition, edited by H. Haken (Springer-Verlag, Berlin, 1979) pp. 210–222.

[28] J. T. Wooton, Nature 413, 841 (2001).

[29] L. A. Maguire and J. W. Porter, Eco Modelling 3, 249 (1977).

[30] A. Okubo and S. A. Levin, Diffusion and Ecological Problems: Modern Perspectives (Springer-Verlag New York, 2001).

[31] M. Scheffer and S. R. Carpenter, TRENDS in Ecology and Evolution 18, 648 (2003).

[32] F. Courchamp, L. Berec, and J. Gascoigne, Allee Effects in Ecology and Conservation (Oxford U Press, 2008).

[33] A. Goyal, V. Dubinkina, and S. Maslov, The ISME Journal 12, 2823–2834 (2018).

[34] V. Dubinkina, Y. Fridman, P. P. Pandey, and S. Maslov, bioRxiv (2018).

[35] C. L. Lehman and D. Tilman, in Spatial ecology: the role of space in population dynamics and interspecific interactions, edited by D. Tilman and P. Kareiva (Princeton U Press, 1997) Chap. 8, pp. 185–203.

