## Supporting Information Appendix for "Spatial Ecology of Territorial Populations"

**This PDF file includes:**

Supplementary text
Figs. S1 to S8
References for SI reference citations

### Supporting Information Text

### 1. Simulation details

**A. Parameters.** In the main text, we often consider distributions of outcomes over multiple random sets of strategies. To choose a random strategy in a competition for two nutrients ($p = 2$), Mathematica's "RandomReal" function was used. To choose a random strategy for $p = 3$, or to choose random initial population abundances, Mathematica's "RandomPoint" method was used to uniformly sample the appropriate simplex.

**B. Well-mixed model.** We compare our spatial territory model to the well-mixed model introduced by Posfai et al. (1). In the well-mixed model, the nutrient concentrations obey

$$\frac{\partial c_i}{\partial t} = S_i - \left(\sum_\sigma n_\sigma \alpha_{\sigma i}\right) r_i(c_i), \quad [1]$$

where $r_i(c_i)$ is an arbitrary uptake function. Nutrient uptake is typically much faster than reproduction, leading to a separation of timescales with $\frac{\partial c_i}{\partial t} = 0$. The population dynamics are then given by

$$\frac{dn_\sigma}{dt} = \left(\sum_i \alpha_{\sigma i} \frac{S_i}{\sum_\gamma n_\gamma \alpha_{\gamma i}} - \delta\right) n_\sigma. \quad [2]$$

**C. Stochastic dynamics.** To simulate the ecological drift shown in Fig. 4B, every $\Delta t$ we generate a "noise" vector $\vec{\nu}$ and set $\vec{n} \to \vec{n} + \vec{\nu}$. The $\sigma$-component of $\vec{\nu}$ is given by $\nu_\sigma = \sqrt{\omega n_\sigma}\xi$, where $\omega$ is the noise strength and $\xi \sim \mathcal{N}(0, 1)$ is a Gaussian random variable. This fixes the variance $\langle \nu_\sigma^2 \rangle = \omega n_\sigma$. If the demographic noise is generated by a Poissonian birth-death process, $\omega = 2\delta\Delta t/\rho$, where $\rho$ is the population density, which fixes the number of individuals as $N_\sigma = \rho n_\sigma$. Since we choose $\Delta t$ and $\omega$ independently, we are effectively varying the population density.

In order to prevent $\vec{\nu}$ from changing the overall population size or leading to negative abundances, we project $\vec{\nu}$ onto the surface $\sum_\sigma \nu_\sigma = 0$ and rescale its magnitude such that $\min(n_\sigma + \nu_\sigma) \geq 0$. In Fig. 4B of the main text, we use noise strength $\omega = 5 \times 10^{-3}$ and period $\Delta t = 5/\delta$, so $\rho = 2000$.

### 2. Fast diffusion expansion

In this section we show that the the well-mixed model is an explicit limit of the spatial territory model. This allows us to make precise comparisons and isolate the effects of spatial structure. The spatial variation of the nutrient environment depends on the nutrient diffusion time $\tau_D \equiv L^2 E/D$. When nutrients diffuse across the whole system much more quickly than species consume them, $\tau_D \to 0$ and the spatial dynamics reduce to the well-mixed dynamics (Eq. 2) with corrections of order $\mathcal{O}(\tau_D)$.

Consider $m$ species competing for $p$ resources. In the region occupied by species $\sigma$, the full equation for the concentration of nutrient $i$ is given by

$$c_{\sigma i}(x) = \frac{S_i}{\alpha_{\sigma i}} + A_{\sigma i} \exp\left(x\sqrt{\frac{\alpha_{\sigma i}}{D}}\right) + B_{\sigma i} \exp\left(-x\sqrt{\frac{\alpha_{\sigma i}}{D}}\right). \quad [3]$$

The spatial coordinate $x$ runs from 0 to $L$, but the nutrient equations are simpler under the change of variables $x \to x - \sum_{\sigma' < \sigma} n_{\sigma'}$, so that each $c_{\sigma i}(x)$ runs from 0 to $n_\sigma$. The following argument applies to each nutrient independently, so we will suppress the nutrient index $i$ for brevity. We can rewrite Eq. 3 in terms of dimensionless variables $u \equiv x/L$, $\tilde{\alpha}_\sigma \equiv \alpha_\sigma/E$, $\tilde{s}_i \equiv S_i/S$, to obtain

$$C_\sigma(u) = \frac{S}{E}\frac{\tilde{s}}{\tilde{\alpha}_\sigma} + A_\sigma \exp\left(u\sqrt{\tilde{\alpha}_\sigma}\epsilon\right) + B_\sigma \exp\left(-u\sqrt{\tilde{\alpha}_\sigma}\epsilon\right), \quad [4]$$

where $\epsilon \equiv \sqrt{\tau_D}$. We will also use relative populations $p_\sigma = n_\sigma/L$, so $\sum_\sigma p_\sigma = 1$. When $\epsilon$ is small, we can expand Eq. 4 around the point $\epsilon = 0$. We require that $C_\sigma(u)$ be continuous and differentiable at the boundaries between populations, to all orders in $\epsilon$. To meet this constraint, we express the coefficients $\{A_\sigma, B_\sigma\}$ as series in $\tau_D$, yielding

$$\begin{aligned} C_\sigma(u) = \frac{S}{E}\frac{\tilde{s}}{\tilde{\alpha}_\sigma} &+ \left(A_\sigma^{(0)} + \epsilon A_\sigma^{(1)} + \epsilon^2 A_\sigma^{(2)} + \ldots\right)\left(1 + \epsilon u\sqrt{\tilde{\alpha}_\sigma} + \epsilon^2 \frac{u^2\tilde{\alpha}_\sigma}{2} + \ldots\right) \\ &+ \left(B_\sigma^{(0)} + \epsilon B_\sigma^{(1)} + \epsilon^2 B_\sigma^{(2)} + \ldots\right)\left(1 - \epsilon u\sqrt{\tilde{\alpha}_\sigma} + \epsilon^2 \frac{u^2\tilde{\alpha}_\sigma}{2} + \ldots\right), \end{aligned} \quad [5]$$

and gather terms of like order to express the nutrient concentration as

$$C_\sigma(u) = \sum_k \epsilon^k C_\sigma^{(k)}(u). \quad [6]$$

It is simpler to use the sum and difference variables $F_\sigma^{(k)} \equiv A_\sigma^{(k)} + B_\sigma^{(k)}$ and $G_\sigma^{(k)} \equiv A_\sigma^{(k)} - B_\sigma^{(k)}$. The $\mathcal{O}(1)$ (or $k = 0$) term is

$$C_\sigma^{(0)} = \frac{S}{E}\frac{\tilde{s}}{\tilde{\alpha}_\sigma} + F_\sigma^{(0)},$$

which is spatially uniform. The only solution to the boundary conditions $C_\sigma^{(0)} = C_{\sigma+1}^{(0)}$ is a constant value across all territories. We call this $C^{(0)}$, and its value will be fixed by the second-order correction.

The terms up to $\mathcal{O}(\epsilon^3)$ are

$$\begin{aligned} C_\sigma^{(1)}(u) &= F_\sigma^{(1)} + u\sqrt{\tilde{\alpha}_\sigma} G_\sigma^{(0)}, \\ C_\sigma^{(2)}(u) &= F_\sigma^{(2)} + u\sqrt{\tilde{\alpha}_\sigma} G_\sigma^{(1)} + \frac{u^2\tilde{\alpha}_\sigma}{2} F_\sigma^{(0)}, \\ C_\sigma^{(3)}(u) &= F_\sigma^{(3)} + u\sqrt{\tilde{\alpha}_\sigma} G_\sigma^{(2)} + \frac{u^2\tilde{\alpha}_\sigma}{2} F_\sigma^{(1)} + \frac{u^3\tilde{\alpha}_\sigma^{3/2}}{6} G_\sigma^{(0)}. \end{aligned} \tag{7}$$

To find the uniform $C^{(0)}$, we write $F_\sigma^{(0)} = C^{(0)} - \frac{S}{E}\frac{\tilde{s}}{\tilde{\alpha}_\sigma}$ and use the requirement that $C_\sigma^{'(2)}(p_\sigma) = C_{\sigma+1}^{'(2)}(0)$ (i.e. the second-order correction be differentiable at the boundaries), yielding

$$\sqrt{\tilde{\alpha}_\sigma} G_\sigma^{(1)} + p_\sigma \tilde{\alpha}_\sigma \left( C^{(0)} - \frac{S}{E}\frac{\tilde{s}}{\tilde{\alpha}_\sigma} \right) = \sqrt{\tilde{\alpha}_{\sigma+1}} G_{\sigma+1}^{(1)}. \tag{8}$$

This gives us an equation at each of the $m$ boundaries. Summing over all $m$ equations, we obtain

$$\begin{aligned} \sum_\gamma \sqrt{\tilde{\alpha}_\gamma} G_\gamma^{(1)} + C^{(0)} \sum_\gamma p_\gamma \tilde{\alpha}_\gamma - \frac{S}{E}\tilde{s} \sum_\gamma p_\gamma &= \sum_\gamma \sqrt{\tilde{\alpha}_\gamma} G_\gamma^{(1)} \\ \therefore C^{(0)} &= \frac{S}{E} \frac{\tilde{s}}{\sum_\gamma p_\gamma \tilde{\alpha}_\gamma}. \end{aligned} \tag{9}$$

Now we want to show that the $\mathcal{O}(\epsilon)$ correction vanishes, i.e. $C_\sigma^{(1)}(u) = 0 \; \forall \; \sigma$. Because $C_\sigma^{(1)}(u)$ is piecewise linear within each territory, the only way to satisfy the boundary conditions of equal $C_\sigma^{(1)}$ and $C_\sigma^{'(1)}$ is $C_\sigma^{(1)} = F_\sigma^{(1)} = constant$, so all the $F_\sigma^{(1)}$ are equal. Then we add up the $m$ equations for $C^{(3)}(u)$ to be differentiable at the boundaries to obtain

$$\begin{aligned} \sum_\gamma \sqrt{\tilde{\alpha}_\gamma} G_\gamma^{(2)} + F^{(1)} \sum_\gamma p_\gamma \alpha_\gamma &= \sum_\gamma \sqrt{\tilde{\alpha}_\gamma} G_\gamma^{(2)} \\ \therefore F^{(1)} &= 0, \end{aligned} \tag{10}$$

so the first-order correction vanishes. So far we have shown (restoring the nutrient index $i$),

$$C_{\sigma i}(u) = \frac{S}{E} \frac{\tilde{s}_i}{\sum_\gamma p_\gamma \tilde{\alpha}_{\gamma i}} + \epsilon^2 C_{\sigma i}^{(2)}(u) + \mathcal{O}(\epsilon^3), \tag{11}$$

where $C_{\sigma i}^{(2)}(u)$ is quadratic in $u$ (Eq. 7) and has coefficients determined by higher-order corrections. Recalling that $\epsilon = \sqrt{\tau_D}$, the population dynamics are then given by

$$\begin{aligned} \frac{dn_\sigma}{dt} &= \sum_i \alpha_{\sigma i} \left( v \int_0^{n_\sigma} C_{\sigma i}(x)\, dx \right) - \delta n_\sigma \\ &= \sum_i \alpha_{\sigma i} \left( v \int_0^{n_\sigma} \frac{S_i}{\sum_\gamma p_\gamma \alpha_{\gamma i}}\, dx + \tau_D v \int_0^{n_\sigma} C_{\sigma i}^{(2)}(x)\, dx \right) - \delta n_\sigma \\ &= \left( \sum_i \alpha_{\sigma i} \frac{v S_i}{\sum_\gamma p_\gamma \alpha_{\gamma i}} - \delta \right) n_\sigma + \mathcal{O}(\tau_D), \\ &= \left( \sum_i \alpha_{\sigma i} \frac{v S_i L}{\sum_\gamma n_\gamma \alpha_{\gamma i}} - \delta \right) n_\sigma + \mathcal{O}(\tau_D). \end{aligned} \tag{12}$$

In units where the nutrient value $v = 1$, the $\mathcal{O}(1)$ term is equivalent to the well-mixed population dynamics (Eq. 2) with a supply vector rescaled by the system size $L$. Therefore, for small $\tau_D$, the spatial dynamics reduce to the well-mixed dynamics with a correction of $\mathcal{O}(\tau_D)$.

### 3. Diffusion time and diversity

In Fig. 2 of the main text, we show that the diffusion time $\tau_D$ is a control parameter for diversity. Why does diversity decrease with increasing $\tau_D$? Figure S1*A* and *B* show the average fraction of surviving species and effective number of species for 500 random communities where $\vec{s} = (0.5, 0.5)$ (*cf.* Fig. 2, where $\vec{s} = (0.4, 0.6)$). Because the nutrient supply is perfectly balanced, there are no oligotrophs to cause extinctions. However, ecosystem diversity still depends on relative abundances. Figure S1*C* shows the average steady-state populations of the three most abundant species from the communities in Fig. S1*A* and *B*.

Around $L = 5$, a transition occurs, and the populations of the Rank-2 and Rank-3 species stop growing with $L$. Instead, the Rank-1 species grows to occupy the additional space. As the population of this dominant species grows, the abundances become less even, reducing biodiversity as measured by both rank-abundance and by effective number of species $M$.

Why does the most abundant species reap all the benefits of increased system size? Consider the example in Fig. S1*D* and *E*, which show the steady-state nutrient concentrations for the same community of species at $L = 20$ and $L = 30$, respectively. When $\tau_D > 1$, the species $\sigma$ with the lowest $R_\sigma = \sum_i S_i/\alpha_{\sigma i}$ will dominate because it has the lowest overall nutrient requirements. ($\sigma$ will drive other species to extinction if it is an oligotroph, but this is not necessary to become the dominant species.) Once $n_\sigma$ is large enough, the $c_{\sigma i}(x) \approx R_{\sigma i}$ in the "bulk" of species $\sigma$'s territory. Recall that $R_{\sigma i}$ is the steady-state nutrient concentration of an isolated population, so $c_i = R_{\sigma i}\ \forall\ i$ gives zero net growth locally. Mathematically, this is expressed as

$$\begin{aligned} \dot{n}_\sigma(x) &\equiv \sum_i \alpha_{\sigma i} c_i(x) - \delta \\ &= \sum_i \alpha_{\sigma i} R_{\sigma i} - \delta \\ &= 0. \end{aligned} \qquad [13]$$

Thus, there is zero local growth in the bulk of the dominant species $\sigma$. As a result, given a steady state $\vec{n}(L)$, a new steady state $\vec{n}(L + \Delta L)$ can always be constructed by inserting $\Delta L$ into the bulk of the dominant species. This change does not affect the boundary fluxes or any other populations, because when $n_\sigma \gg \sqrt{E/D}$, the concentration $c_\sigma(x)$ can be approximated as a uniform bulk with two boundary layers that are insensitive to the size of the bulk.

Typically, *only* the most dominant species $\sigma$ will have such a bulk region. Even communities with very similar strategies have a single dominant species: in Fig. S1*D* and *E*, $\alpha_{\sigma 1} - \alpha_{\sigma' 1} < 0.014$, but $\sigma$ is clearly dominant. As a result, inserting $\Delta L$ into $n_{\sigma'}$ would not yield a steady state, because this species has no bulk region where $\dot{n}_{\sigma'}(x) = 0$. The spatial model exhibits alternative steady states, so it is possible that the expanded system reaches other steady states than $n_\sigma \to n_\sigma + \Delta L$. However, Fig. S1*C* suggests that the new steady state always has a dominant species which absorbs the $\Delta L$, even if the identity of that species can change with system size.

Finally, it is worth noting that at high $\tau_D$, some sets of strategies lead to a computationally stiff system. In Fig. 2*C* and *D* and Fig. S1*A* and *B*, we analyzed each set of strategies across the full range of $\tau_D$. When Mathematica detected a stiff system at high $\tau_D$, we excluded those competitions from analysis at any $\tau_D$. For example, the point $\tau_D = 10$ only includes sets of strategies that did not lead to computational stiffness at high $\tau_D$. (We used 400 sets of species satisfying this criterion in Fig. 2 in the main text.) To check that this filter did not bias our results, we confirmed that the distribution of strategies present in our data was uniform (Fig. S1*F*). We then compared the diversity statistics $P(M)$ of our filtered data for $\tau_D = 10$ (Fig. S1*G*) to 2000 competitions with the same conditions but no filtering (Fig. S1*H*), and observed similar distributions. The Kolmogorov-Smirnov test, as implemented by Mathematica's "KolmogorovSmirnovTest," returned a $p = 0.78$, consistent with data coming from the same underlying distribution. Thus the conclusions drawn from Fig. S1*A* and *B* (along with Fig. 2 of the main text) are unlikely to be affected by the filtering process.

### 4. Rank abundance curves

Throughout the text, we have characterized steady-state diversity using the fraction of surviving species and the effective number of species. However, ecologists often employ rank abundance curves to visualize the full distribution of abundances in an ecosystem. Fig. S2 shows steady-state rank abundance curves for unbalanced and balanced nutrient supplies across a range of $\tau_D$. Each point is an average over many random communities. For both nutrient supplies, the well-mixed system ($\tau_D = 0$) has a shallow curve, indicating that all species have similar abundances. (This is for the case of equal initial abundances. Because the well-mixed system has a degenerate manifold of steady states, less even initial conditions typically yield less even distributions.) When the nutrient supply is unbalanced (left), even rapid nutrient mixing ($\tau_D = 0.01$) allows the most abundant species to occupy $\approx 70\%$ of space. As a result, the other species' shares of abundances decrease precipitously with rank. Increasing $\tau_D$ exacerbates this effect, with the rank-1 species controlling $> 90\%$ of space at $\tau_D = 1600$. It should be noted that most steady states with an unbalanced nutrient supply have fewer than ten surviving species, so many abundances contributing to the high rank averages are zero. A balanced nutrient supply leads to higher diversity (right), but the population of the rank-1 species still grows from $50 - 85\%$ as the nutrient mixing time increases.

### 5. Linear stability analysis and slow directions

When there are more species $m$ than nutrients $p$, the spatial model has a slow manifold of dimension $m - p$ in population space, on which the relaxation to steady state takes place on a timescale $t_{\text{slow}} \sim 1/\tau_D$. Specifically, linear stability analysis around the fixed point yields $m - p$ eigenvalues $\lambda_i \propto -\tau_D$. Figure S3 shows an example for $(m, p) = (3, 2)$. Fig. S3A shows the directions of the eigenvectors $\vec{v}_i$ in population space. The first eigenvector $\vec{v}_1$ is the "stiff" mode of changing $n_1$, and has a large eigenvalue that does not scale with $\tau_D$ (Fig. S3B). $\vec{v}_2$ is the "soft mode" of the balance between $n_2$ and $n_3$. This is the slow manifold, with $|\lambda_2| \propto \tau_D$ (Fig. S3C). As $\tau_D \to 0$ in the well-mixed limit, $\vec{v}_2$ becomes the degenerate manifold of well-mixed steady states. $\vec{v}_3$ corresponds to perturbations that change the total population size $L$. Because the dynamics do not allow $\vec{n}$ to leave the surface $\sum_\sigma n_\sigma = L$, this is a new system so $\lambda_3$ is not relevant to the dynamics.

Below, we show that the slow manifold is a general feature of the spatial model. To do so, we calculate the Jacobian, analyze the stability of the well-mixed dynamics, and treat the spatial dynamics as a perturbation of the well-mixed eigenvalues.

**A. Calculating the Jacobian.** We found in Section 2 that we can express the population dynamics as a series in $\tau_D$ by expanding $c(x)$:

$$\frac{dn_\sigma}{dt} = \left(\sum_i \alpha_{\sigma i} \frac{S_i L}{\sum_\mu n_\mu \alpha_{\mu i}} - \delta\right) n_\sigma + \tau_D f_\sigma(\vec{n}), \quad [14]$$

where $f_\sigma(\vec{n}) \equiv \sum_i \alpha_{\sigma i} \int c^{(2)}_{\sigma i}(x)dx$ and we have truncated the series at $\mathcal{O}(\tau_D)$.

At steady state,

$$\left(\sum_i \alpha_{\sigma i} \frac{S_i L}{\sum_\mu n^*_\mu \alpha_{\mu i}} - \delta\right) n^*_\sigma + \tau_D f_\sigma(\vec{n}^*) = 0, \quad [15]$$

which implies that in region $\sigma$,

$$\frac{S_i L}{\sum_\mu n^*_\mu \alpha_{\mu i}} = \frac{\delta}{E} + \tau_D \frac{Q_{\sigma i}(\vec{n}^*)}{\alpha_{\sigma i}}, \quad [16]$$

where $Q_{\sigma i}$ satisfies

$$n^*_\sigma \sum_i Q_{\sigma i}(\vec{n}^*) = -f_\sigma(\vec{n}^*). \quad [17]$$

Eq. 16 says that at steady state, the nutrient concentrations $c_{\sigma i} = \delta/E$ at $\mathcal{O}(1)$ for all $i$ and $\sigma$, with spatial corrections of $\mathcal{O}(\tau_D)$ which depend on both the nutrient and the region. (Note that the total population is fixed, so $\delta = S$ in units where $v = 1$. Thus this is equivalent to $c_{\sigma i} = S/E$.) This simplifies calculation of the matrix elements of the Jacobian $J$:

$$\left.\frac{\partial \dot{n}_\sigma}{\partial n_\gamma}\right|_{\vec{n}^*} = -\sum_i \frac{\alpha_{\sigma i}\alpha_{\gamma i}}{S_i L}\left(\frac{S_i L}{\sum_\mu n^*_\mu \alpha_{\mu i}}\right)^2 n^*_\sigma + \tau_D \partial_{n_\gamma} f_\sigma(\vec{n}^*) \quad [18]$$

$$= -\sum_i \frac{\alpha_{\sigma i}\alpha_{\gamma i}}{S_i L}\left(\frac{\delta}{E} + \tau_D^2 \frac{Q_{\sigma i}(\vec{n}^*)}{\alpha_{\sigma i}}\right)^2 n^*_\sigma + \tau_D \partial_{n_\gamma} f_\sigma(\vec{n}^*) \quad [19]$$

$$= -\sum_i \frac{\alpha_{\sigma i}\alpha_{\gamma i}}{S_i L}\left(\frac{\delta}{E}\right)^2 n^*_\sigma + \tau_D g_{\sigma\gamma}(\vec{n}^*), \quad [20]$$

where $g_{\sigma\gamma}(\vec{n}^*)$ is a complicated function containing all the spatial interactions. This expression holds for the diagonal elements $\sigma = \gamma$ as well. Now we can express $J$ as the well-mixed Jacobian with an $\mathcal{O}(\tau_D)$ correction:

$$\begin{aligned} J &= -\left(\frac{\delta}{E}\right) DM + \tau_D G \\ J &= J_0 + \tau_D G, \end{aligned} \quad [21]$$

where we have used the fact that $L = \sum_\sigma n_\sigma = \delta/E$. In Eq. 21, $J_0 \equiv -(\frac{\delta}{E})DM$ is the Jacobian of the well-mixed system, $M_{\sigma\gamma} = \sum_i \frac{\alpha_{\sigma i}\alpha_{\gamma i}}{S_i}$, $G_{\sigma\gamma} = g_{\sigma\gamma}(\vec{n}^*)$, and

$$D = \begin{pmatrix} n^*_1 & 0 & \cdots & 0 \\ 0 & n^*_2 & \cdots & 0 \\ \vdots & \vdots & & \vdots \\ 0 & 0 & \cdots & n^*_m \end{pmatrix}.$$

**B. Well-mixed stability analysis.** Before studying the effect of the spatial perturbation, we first determine the stability of the unperturbed (well-mixed) system. $J_0$ has $m$ eigenvalues $\lambda_j$, and below we show that for $m - p$ of these, $\lambda_j = 0$. These zero eigenvalues correspond to the $m - p$ dimensional degeneracy of steady states.

To find the number of zero eigenvalues, it suffices to find the number of linearly independent solutions to the equation

$$DM\mathbf{x} = \mathbf{0}. \quad [22]$$

Writing this in terms of matrix elements, we have

$$n^*_\sigma \sum_i \alpha_{\sigma i} y_i = 0, \quad [23]$$

where $y_i \equiv (\alpha_{1i}x_1 + \alpha_{2i}x_2 + ... + \alpha_{mi}x_m)/S_i$. This is a system of $m$ equations with $p$ unknowns, where $m > p$. The number of independent equations is the number of linearly independent strategies. Thus, if there are at least $p$ linearly independent strategies, the only solution is $y_i = 0 \;\forall\; i$.

Then the system $y_i = \alpha_{1i}x_1 + \alpha_{2i}x_2 + ... + \alpha_{mi}x_m = 0$ has $m$ unknowns but only $p$ equations, so there are $m - p$ linearly independent solutions in $\mathbf{x}$. Thus the Jacobian $J_0$ of the well-mixed system has $m - p$ zero eigenvalues.

**C. Spatial perturbation.** Now we show that the spatial dynamics encoded in $\tau_D G$ in Eq. 21 modify the zero eigenvalues of the well-mixed Jacobian $J_0$, transforming the degenerate manifold of steady states into a slow manifold with a timescale $t_{\text{slow}} \sim 1/\tau_D$.

Consider the matrix $A+\epsilon B$, where $\epsilon \ll 1$. Assume the matrix $A \in \mathbb{R}^{m\times m}$ is diagonalizable, that it has $p$ non-zero eigenvalues $\lambda_1, ..., \lambda_p$, and that it has $m-p$ zero eigenvalues. Because $A$ is diagonalizable, it can be decomposed into $A = V\Lambda V^{-1}$, where

$$\Lambda = \left(\begin{array}{c|c} \Lambda_{11} & 0 \\ \hline 0 & 0 \end{array}\right), \tag{24}$$

and $\Lambda_{11}$ is the $p\times p$ diagonal matrix with entries $\lambda_1, ...\lambda_p$. We look for the small eigenvalues of $A+\epsilon B$ in the form $\lambda = \epsilon c$. The characteristic polynomial of $A+\epsilon B$ is given by

$$\begin{aligned} f(A+\epsilon B) &= \det\left(\Lambda + \epsilon V^{-1}BV - \epsilon c\mathbf{I}\right) \\ &= \det\left(\Lambda + \epsilon(B' - c\mathbf{I})\right) \end{aligned} \tag{25}$$

where $B' \equiv V^{-1}BV$. We write $B'$ in terms of block matrices:

$$B' = \left(\begin{array}{c|c} B'_{11} & B'_{12} \\ \hline B'_{21} & B'_{22} \end{array}\right) \to \Lambda + \epsilon(B' - c\mathbf{I}) = \left(\begin{array}{c|c} \Lambda_{11} + \epsilon(B'_{11} - c\mathbf{I}) & \epsilon B'_{12} \\ \hline \epsilon B'_{21} & \epsilon(B'_{22} - c\mathbf{I}) \end{array}\right). \tag{26}$$

For a block matrix $M = \left(\begin{array}{c|c} M_{11} & M_{12} \\ \hline M_{21} & M_{22} \end{array}\right)$, $\det(M) = \det(M_{11})\cdot\det(M_{22} - M_{21}M_{11}^{-1}M_{12})$ if $M_{11}$ is invertible. $\Lambda_{11} + \epsilon(B'_{11} - c\mathbf{I})$ is invertible for small $\epsilon$, so the characteristic polynomial from Eq. 25 can be expressed as

$$\begin{aligned} f(A+\epsilon B) &= \det\left(\Lambda_{11} + \epsilon(B'_{11} - c\mathbf{I})\right)\cdot\det\left(\epsilon(B'_{22} - c\mathbf{I}) - \epsilon^2 B'_{21}\left(\Lambda_{11} + \epsilon(B'_{11} - c\mathbf{I})\right)^{-1}B'_{12}\right) \\ &= \det\left(\Lambda_{11} + \epsilon(B'_{11} - c\mathbf{I})\right)\cdot\epsilon^{m-p}\cdot\det\left((B'_{22} - c\mathbf{I}) - \epsilon B'_{21}\left(\Lambda_{11} + \epsilon(B'_{11} - c\mathbf{I})\right)^{-1}B'_{12}\right) \\ &= \left(\prod_{i=1}^{p}\lambda_i + \mathcal{O}(\epsilon)\right)\cdot\epsilon^{m-p}\cdot\left(\det(B'_{22} - c\mathbf{I}) + \mathcal{O}(\epsilon)\right) \end{aligned} \tag{27}$$

The small eigenvalues $\epsilon c$ can be found by dropping terms of $\mathcal{O}(\epsilon)$ and solving for $c$:

$$\left(\prod_{i=1}^{p}\lambda_i\right)\cdot\det(B'_{22} - c\mathbf{I}) = 0, \tag{28}$$

where $c$ is independent of $\epsilon$. Thus the matrix $A+\epsilon B$ has $m-p$ eigenvalues of the form $\lambda = \epsilon c$, corresponding the $m-p$ solutions of Eq. 28.

Therefore the Jacobian $J_0 + \tau_D G$, has $m-p$ eigenvalues $|\lambda| \propto \tau_D$, just as we found via numerical stability analysis. The slow manifold is a general feature of our spatial model whenever there is a diverse community with more species than nutrients ($m > p$).

### 6. Alternative steady states

Figure 5 of the main text shows that spatial structure can lead to multistability and the Allee effect. Figure S4 classifies the steady states for the full space of strategies when two species compete for two nutrients. In the well-mixed model, this competition would always have a single steady state because the number of species equals the number of nutrients. Now, however, similar initial populations can produce dramatically different outcomes. In the blue regions in Fig. S4 labeled "Competitive exclusion", both species specialize strongly in the same resource, and there is a single steady state in which one species drives the other to extinction. In the brown regions labeled "Coexistence", the two species specialize in different resources, and there is a single steady state in which they coexist. In the tan region, both species specialize in Nutrient 2 but have strategies similar to the nutrient supply vector, and we observe the Allee effect: Species 2 comprises half or more of the system if its initial population passes a threshold, but goes extinct otherwise. Finally, in the beige region labeled "Bistable coexistence", there are two steady states where both species coexist. Typically, one of these steady states will have relatively equal populations, but not the other, (see Fig. 5), so a small initial variation can change the final community composition substantially.

The fraction of parameter space in which alternative steady states occur depends on the nutrient environment. Figure S4*B* and *D* show that making the nutrient supply more symmetric shrinks the regions of multistability and the Allee effect. On the other hand, Fig. S4*C* and *D* show that increasing the diffusion time $\tau_D$ substantially increases the size of these regions. Thus $\tau_D$ not only controls the diversity of steady states (Fig. 2, main text and Fig. S1), but also determines whether alternative outcomes are possible.

### 7. Unequal enzyme budgets

Figure 6 of the main text demonstrates that in spatial ecosystems diversity beyond the competitive-exclusion limit persists for unequal enzyme budgets. In fact, diversity peaks for intermediate variance in the enzyme budget $E$. Figure S5 shows that this is due to an asymmetric effect on oligotroph versus non-oligotroph strategies. In Fig. S5*A*, $\delta E^*$ is the *decrease* in $E$ at which an oligotroph stops driving others extinct, or the *increase* in $E$ at which a non-oligotroph strategy begins driving others extinct. In other words, $\delta E^*$ is how much the enzyme budget must change for an oligotroph to behave as a non-oligotroph, or vice versa. The further a strategy is from the oligotroph region, the more $E$ must increase for that species to drive others extinct like an oligotroph. In the oligotroph region, however, a very small decrease in $E$ allows the oligotroph to coexist with many other species. As a result, the random enzyme budgets in Fig. 6 often suppress the oligotroph effect without allowing other species to become dominant.

Why are oligotrophs so sensitive to changes in enzyme budget? Recall that oligotrophs are defined by $R_\sigma < p$. Figure S5*B* shows $R_\sigma$ as a function of strategy. It is suggestive that in the oligotroph region, $R_\sigma$ is very close to the threshold $p$, whereas strategies away from the oligotroph region have $R_\sigma \gg p$. For any particular species and community, the critical budget change $\delta E^*$ is not simply the change in $E$ necessary to cross $p$, but some more complicated function of $\tau_D$ and the other species present. However, Fig. S5*C* shows that $\delta E^*$ increases monotonically with $R_\sigma$. In particular, it rapidly increases as $R_\sigma$ crosses the oligotroph threshold before leveling off. At lower $\tau_D$ there is more variation (Fig. S5*D*), but the separate scales for oligotrophs and non-oligotrophs remain apparent.

### 8. Lattice territory model

We have shown that spatial territories reduce biodiversity in a one-dimensional model. Does this remains true in two spatial dimensions? In 2D, each species will have more neighbors, and there are more opportunities for pattern formation. However, implementing our model in a 2D continuous space presents serious challenges. In 1D, territories simply expand or contract as populations change. In 2D, there are many possible ways for territories to grow, depending on the details of mechanical forces acting at boundaries between populations. (For instance, one might model the boundary as discrete individuals shoving each other, or as a fluid interface subject to fingering instabilities, etc.) To bypass these complications, we propose a lattice territory model which captures the two essential features of the continuous model: non-overlapping territories and a nutrient environment globally coupled via diffusion. In this section, we show that the 1D version of this lattice territory model qualitatively reproduces diversity patterns from the 1D continuous model. Then we demonstrate that our conclusions regarding biodiversity hold in 2D versions of the lattice territory model, indicating that our primary results are not unique to one dimension.

**A. The model.** We consider $m$ species competing for $p$ resources on a lattice with $m$ sites. Each lattice site constitutes a "territory" occupied by a single species $\sigma$. A species never leaves its territory, but can exchange nutrients with its neighbors on the lattice (see Fig. S6*A* for an illustration). Within each territory, the nutrients are assumed to be well-mixed. The population size $n_\sigma$ in each territory does not influence the spatial arrangement of territories on the lattice, but does influence the flux of nutrients between territories. The total amount of nutrient $i$ in territory $\sigma$, $N_{\sigma i}$, obeys

$$\frac{\partial N_{\sigma i}}{\partial t} = S_i n_\sigma - \alpha_{\sigma i} n_\sigma c_{\sigma i} + D \sum_{\sigma'} \left( c_{\sigma' i} - c_{\sigma i} \right), \qquad [29]$$

where $c_{\sigma i} = N_{\sigma i}/n_\sigma$ is the concentration of nutrient $i$ in territory $\sigma$. The sum over $\sigma'$ includes all $z$ neighbors of species $\sigma$ on the lattice. Note that the flux of each nutrient into a territory from the external supply scales with the territory size $n_\sigma$, as in our continuous 1D model. If nutrient levels equilibrate much more quickly than populations change, we can again assume a separation of timescales ($\partial N/\partial t = 0$) and obtain equations for the steady-state nutrient concentrations:

$$c_{\sigma i} = \frac{S_i n_\sigma + D \sum_{\sigma'} c_{\sigma' i}}{\alpha_{\sigma i} n_\sigma + Dz}. \qquad [30]$$

Note that if $D = 0$, $c_{\sigma i} = S_i/\alpha_{\sigma i} = R_{\sigma i}$, which is the nutrient environment of an isolated species in the 1D continuous model. In the case of an extinction, with $n_\sigma = 0$, we note that $c_{\sigma i} = \sum_{\sigma'} c_{\sigma' i}/z$, so nutrients still flow through the territories of extinct species. Because the nutrients are well-mixed within a territory, the population dynamics are simply given by

$$\frac{dn_\sigma}{dt} = \left( \sum_i \alpha_{\sigma i} c_{\sigma i} - \delta \right) n_\sigma. \qquad [31]$$

The total system population size $\sum_\sigma n_\sigma = n_T$ remains fixed so long as $S = \delta$. As in the continuous 1D model, we assume exact metabolic trade-offs, i.e. $\sum_i \alpha_{\sigma i} = E$ for all species, as well as periodic boundary conditions.

**B. Lattice diffusion time.** In the 1D continuous model, we define the nutrient mixing timescale as the time to diffuse across the system relative to the uptake time: $\tau_D = \frac{L^2/D}{1/E} = \frac{L^2 E}{D}$. This can also be expressed in terms of the system size and the decay length $\lambda$, the typical distance nutrients diffuse before being consumed: $\tau_D = (\frac{L}{\lambda})^2 = (\frac{L}{\sqrt{D/E}})^2$. We will use this second definition to calculate $\tau_D$ in the lattice territory model.

In the lattice territory model, the system size $L$ will be replaced by the number of sites in one spatial dimension. We calculate the decay length $\lambda$ by considering a 1D lattice with a source at the leftmost site. (Here, we do not use periodic boundary conditions.) At each site, nutrients can either diffuse to the next site or be consumed. How does the nutrient concentration decay as a function of distance from the source?

Specifically, we consider a 1D lattice of sites $\sigma = 1...m$, with a nutrient supply $s$ at site $\sigma = 1$. For simplicity, we assume uniform populations and strategies ($n_\sigma = n$ and $\alpha_\sigma = \alpha$) at all sites. We wish to calculate the steady-state nutrient concentrations $c(\sigma)$. These are given by

$$\begin{aligned} s - \alpha c(1) + \frac{D}{n}\Big(c(2) - c(1)\Big) &= 0, \\ -\alpha c(m) + \frac{D}{n}\Big(c(m-1) - c(m)\Big) &= 0, \\ -\alpha c(\sigma) + \frac{D}{n}\Big(c(\sigma+1) + c(\sigma-1) - 2c(\sigma)\Big) &= 0, \quad 1 < \sigma < m. \end{aligned} \tag{32}$$

Defining $x \equiv \alpha\frac{n}{D} + 2$, we can rearrange these equations to yield

$$\begin{aligned} c(1) &= \frac{s\frac{n}{D} + c(2)}{x-1}, \\ c(m) &= \frac{c(m-1)}{x-1}, \\ c(\sigma) &= \frac{c(\sigma+1) + c(\sigma-1)}{x}, \quad 1 < \sigma < m. \end{aligned} \tag{33}$$

In general, the solution to this system of $m$ equations is quite complicated. We proceed by finding the ratios of the concentrations between adjacent sites, then studying the limit $m \gg 1$, which yields an exponential form for $c(\sigma)$. From Eq. 33, we see

$$\frac{c(m)}{c(m-1)} = \frac{1}{x-1}. \tag{34}$$

$1/(x-1) < 1$, so the last site has a lower concentration than the penultimate site, as expected. We can find the subsequent concentration ratios iteratively:

$$\begin{aligned} \frac{c(m-1)}{c(m-2)} &= \left(x - \frac{1}{x-1}\right)^{-1}, \\ &\cdots \\ \frac{c(\sigma)}{c(\sigma-1)} &= f(m-\sigma), \quad \sigma > 2. \end{aligned} \tag{35}$$

The proportionality constant $f(k)$ is defined iteratively:

$$\begin{aligned} f(k) &= \Big(x - f(k-1)\Big)^{-1}, \\ f(0) &= \frac{1}{x-1}. \end{aligned} \tag{36}$$

The recurrence relation Eq. 35 does not hold for $c(2)/c(1)$ because of the source at $\sigma = 1$. Using Eq. 33 and Eq. 35, we find the following solution for the nutrient profile:

$$\begin{aligned} c(2) &= (x-1)c(1) - s\frac{n}{D}, \\ c(3) &= c(2)f(m-2), \\ &\cdots \\ c(\sigma) &= \left((x-1)c(1) - s\frac{n}{D}\right)\prod_{l=2}^{\sigma-1} f(m-l). \end{aligned} \tag{37}$$

We seek only the functional dependence of $c(\sigma)$ on $\sigma$, so we can leave the boundary term $c(1)$ unevaluated. Equation 37 is exact, but we are interested in the decay of $c(\sigma)$ near the source, in the limit of many sites. Then $m - l >> 1$ for every term in the product, and we can focus on the behavior of $f(k)$ as $k$ becomes large. Note that $f(k)$ can be written as a continued fraction:

$$
\begin{aligned}
f(1) &= \frac{1}{x - \frac{1}{x-1}}, \\
f(2) &= \frac{1}{x - \frac{1}{x - \frac{1}{x-1}}}, \\
f(3) &= \frac{1}{x - \frac{1}{x - \frac{1}{x - \frac{1}{x - \frac{1}{x-1}}}}}, \\
&\dots
\end{aligned}
\qquad [38]
$$

Consider the limit $F \equiv \lim_{k\to\infty} f(k)$. An infinite continued fraction converges if the sequence of convergents approaches a limit. $f(k)$ satisfies this property for $x \geq 2$ (i.e. $\alpha n/D \geq 0$), so the limit $F$ exists. Then we can solve the following equation for $F$:

$$
\begin{aligned}
F &= \frac{1}{x - F}, \\
\rightarrow F &= \frac{x \pm \sqrt{x^2 - 4}}{2}.
\end{aligned}
\qquad [39]
$$

We choose the smaller of the two solutions on the physical grounds that $F < 1$, so each site has a smaller concentration than the last. Then if $m \gg \sigma$, we can approximate Eq. 37 as

$$
\begin{aligned}
c(\sigma) &\approx \left( (x-1)c(1) - s\frac{n}{D} \right) \prod_{l=2}^{\sigma-1} F \\
&= \left( (x-1)c(1) - s\frac{n}{D} \right) F^{\sigma-2} \\
&= \left( (x-1)c(1) - s\frac{n}{D} \right) \exp(-2\log F)\exp(-\sigma/\lambda),
\end{aligned}
\qquad [40]
$$

where

$$
\lambda \equiv -\frac{1}{\log F}. \qquad [41]
$$

We have achieved our goal of calculating the decay length $\lambda$. However, we are interested in comparing the lattice territory model to the 1D continuous model, where the entire system is diffusively coupled. We expect that the lattice model will be most directly relevant to the continuous model when $\lambda$ spans at least a few sites. This suggests taking the limit $\alpha n/D \ll 1$. Then we can approximate $F \approx 1 - \sqrt{\alpha n/D}$, where we have discarded terms of order $\alpha n/D$ or higher, and expand the logarithm to obtain

$$
\lambda \approx \frac{1}{\sqrt{\alpha n/D}}. \qquad [42]
$$

This result holds for the simplified model with a point source, but we can use it to characterize the diffusion time in the full lattice territory model by considering the geometry. In the 1D lattice territory model, the system size $L$ is the number of sites $m$. The typical scales of $\alpha$ and $n$ are set by $E$ and $n_T/m$, so

$$
\tau_D^{d=1} = \left(\frac{L}{\lambda}\right)^2 = m\frac{n_T E}{D}. \qquad [43]
$$

In the 2D lattice territory model, the linear dimensions of the system are given by the number of sites per row and number of sites per column. We consider lattices of $m$ sites with an equal number of rows and columns, so there will be a single linear dimension $L = \sqrt{m}$. The typical scale of population sizes will again be $n_T/m$. This gives

$$
\tau_D^{d=2} = \left(\frac{L}{\lambda}\right)^2 = \frac{n_T E}{D}. \qquad [44]
$$

For the rest of Section 8 and in Fig. S6-S8, Eq. 43 and Eq. 44 define $\tau_D$ when we use it with respect to a particular lattice territory model. When referring to the continuous model, we use the main text's definition with $L = n_T$, so $\tau_D = n_T^2 E/D$.

**C. 1D lattice.** How do outcomes in the 1D lattice model compare to outcomes in the 1D continuous model of the main text? Figure S6*B* shows the outcome of 16 species competing for two resources. The left subplot uses the continuous model, and the right subplot uses the lattice model. In the well-mixed case, all 16 species would coexist. We see that both spatial models allow coexistence beyond the competitive-exclusion limit, but much of the biodiversity is lost relative to the well-mixed case. Indeed, even the identity of the surviving species is similar between the two 1D models, although the outcome in the lattice model is slightly more diverse. Figure S6*C* shows how the effective number of species $M$ at steady state depends on the nutrient diffusion time $\tau_D$, for the same strategies and nutrient supply as $A$. In both models, increasing $\tau_D$ reduces biodiversity by making the abundances of survivors less equal. Finally, Fig. S6*D* shows the outcome of a competition with the same nutrient supply and strategies as $A$, except that the oligotroph strategy $\vec{\alpha}_4$ has been moved outside the oligotroph region. In both models, the number of species coexisting at steady state increases dramatically. Thus the 1D lattice territory model recapitulates three essential features of the 1D continuous model:

- Spatial structure reduces biodiversity relative to the well-mixed case, but allows coexistence beyond competitive exclusion.
- $\tau_D$ acts as a control parameter for biodiversity by decreasing the evenness of abundances.
- The presence of oligotroph strategies leads to low-diversity steady states.

Moreover, the lattice territory model generalizes readily to 2D, allowing us to ask how dimensionality might affect these features of the model.

**D. 2D lattices.** How does the extension of the lattice territory model into two spatial dimensions affect biodiversity? Figure S7*A* shows the outcome of competition for the same strategies and supply as Fig. S6*B* and *C*, but with competitors arranged on square and hexagonal lattices. These situations differ from each other and the 1D case in their connectivity. As an example, consider Species 6. In 1D, Species 6 directly exchanges nutrients with two neighbors ($\sigma' \in \{5, 7\}$). However, the number of neighbors increases to four on the square lattice ($\sigma' \in \{2, 5, 7, 10\}$) and to six on the hexagonal lattice ($\sigma' \in \{2, 3, 5, 7, 10, 11\}$). In spite of these differences, the impact of spatial structure on biodiversity remains consistent: diversity is lost compared to the well-mixed model, but exceeds the competitive-exclusion limit. (There are intriguing differences in the number, strategies, and spatial arrangements of surviving species, which will be explored in future studies.)

Figure S7*B* shows how the 2D communities in *A* are affected by the nutrient diffusion time $\tau_D$. Again, increasing $\tau_D$ reduces the effective number of species by rendering abundances less even. Figure S7C shows the outcomes of competition for the same community as Fig. S7*A* and *B*, except with the oligotroph stategy $\vec{\alpha}_4$ moved outside the oligotroph region. Again, diversity increases dramatically in the absence of oligotrophs.

In Fig. S8 we show that these patterns are not particular to the strategies in Fig. S6 and S7, but are generic features of the lattice model. Figure S8*A* shows the mean fraction of species surviving at steady state for 300 random sets of 16 strategies, across all three geometries, with and without oligotrophs. For every lattice, spatial structure reduces diversity relative to the well-mixed case, and removing oligotrophs allows many more species to coexist. Indeed, the oligotroph effect seems to be amplified for 2D lattices. Figure S8*B* shows the effective number of species at steady state as a function of $\tau_D$, averaged over many random communities. $\tau_D$ controls the evenness of abundances in all cases, just as it does in the 1D continuous model (*cf.* Fig. 2D in the main text).

There are many interesting features of the lattice model which deserve further investigation. For now, however, we have demonstrated that it captures the essential biodiversity patterns of the 1D continuous model, and shown that these generalize to two spatial dimensions. This suggests that our conclusions from the main text – namely, that spatial structure can reduce biodiversity in a resource-competition model with metabolic trade-offs – are not restricted to 1D, nor are they particularly sensitive to model details.

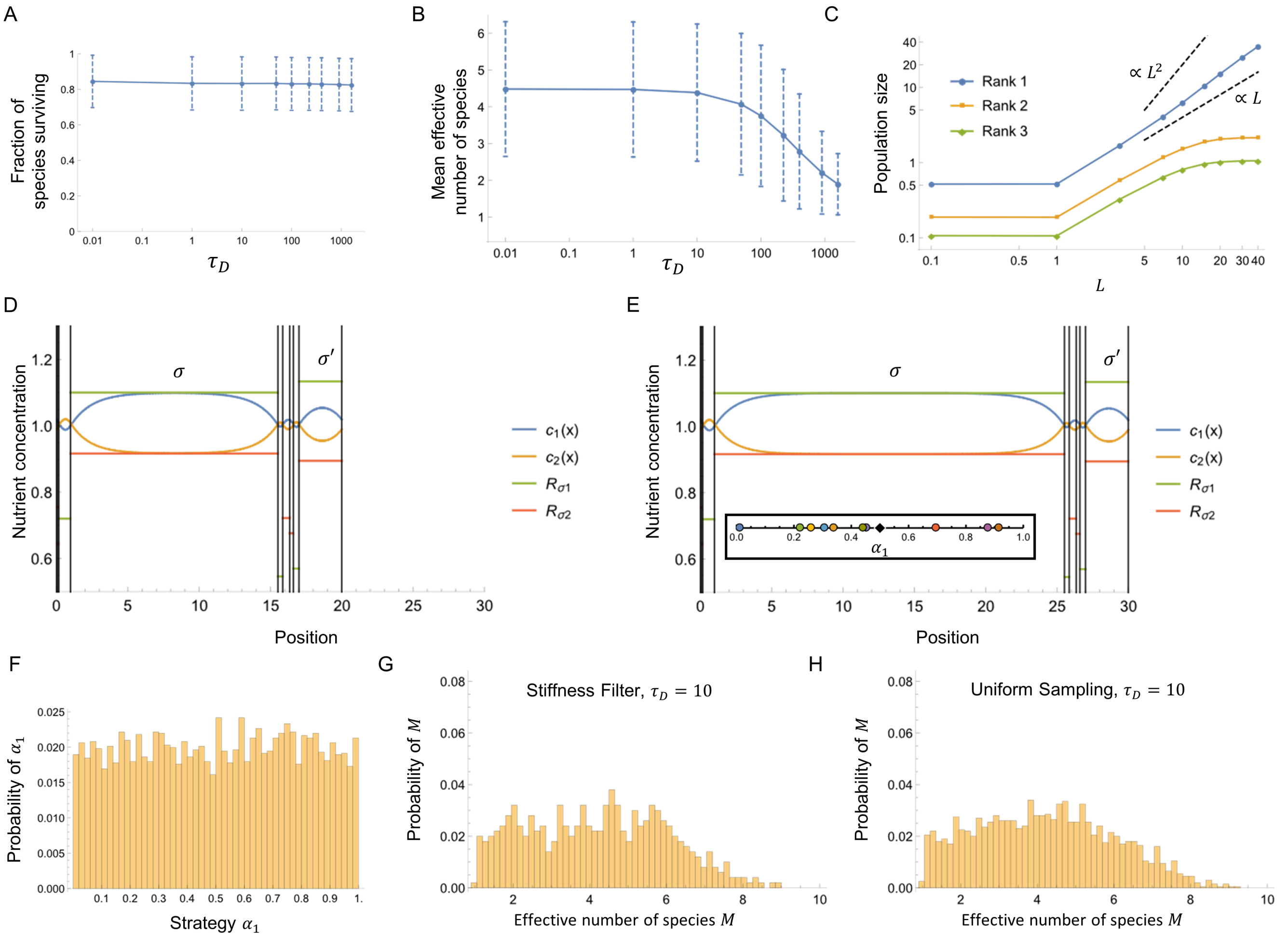


**Fig. S1.** $\tau_D$ controls diversity by setting the abundance of the most dominant species. (*A*) Fraction of initial species coexisting at steady state with $\vec{s} = (0.5, 0.5)$ at various $\tau_D$. A population is considered extinct if $n_\sigma/L < 10^{-6}$ (mean $\pm$ SD for 500 random sets of ten strategies). (*B*) Effective number of species $M$ at steady state (mean $\pm$ SD for same strategies as *A*). (*C*) System size $L$-dependence of population size for the three most abundant species (Rank 1, 2, and 3), averaged over 500 randomly-selected sets of 10 species. At large $L$, only the dominant species grows with increasing system size. (*D*) Steady-state nutrient concentrations for an example community with $L = 20$. Species territories are separated by vertical black lines, and each region shows the nutrient concentrations $R_{\sigma i}$ which the local species would maintain in isolation. (*E*) Same as *D*, but with $L = 30$. (Inset) Strategies and nutrient supply for *D* and *E*. (*F*) Distribution of strategies present in the 500 random communities of *A-C*, which passed a test for numerical stiffness at high $\tau_D$. The distribution is uniform. (*G*) Full distribution of effective number of species $M$ for the communities in *A-C* at $\tau_D = 10$. (*H*) Distribution of effective number of species $M$ for 2000 communities at $\tau_D = 10$, with no stiffness test. The Kolmogorov-Smirnov test on *G* and *H* yields a p-value of $0.78$, consistent with the data come from the same underlying distribution.

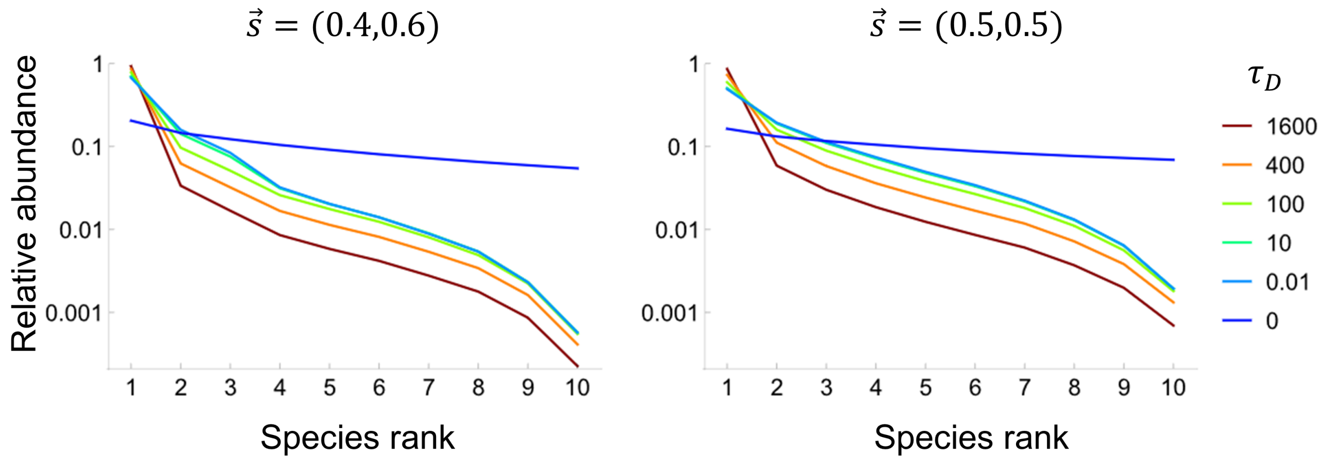


**Fig. S2.** Rank abundance curves show that increasing the nutrient diffusion time $\tau_D$ reduces the evenness of abundances. *Left:* Steady-state rank-abundance curves for the unbalanced nutrient supply $\vec{s} = (0.4, 0.6)$, across a range of $\tau_D$. Each curve is averaged over 400 randomly-selected sets of ten initial strategies, and each simulation begins with equal initial populations. *Right:* Steady-state rank-abundance curves for the balanced nutrient supply $\vec{s} = (0.5, 0.5)$, across a range of $\tau_D$. Each curve is averaged over 500 randomly-selected sets of ten initial strategies, and each simulation begins with equal initial populations.

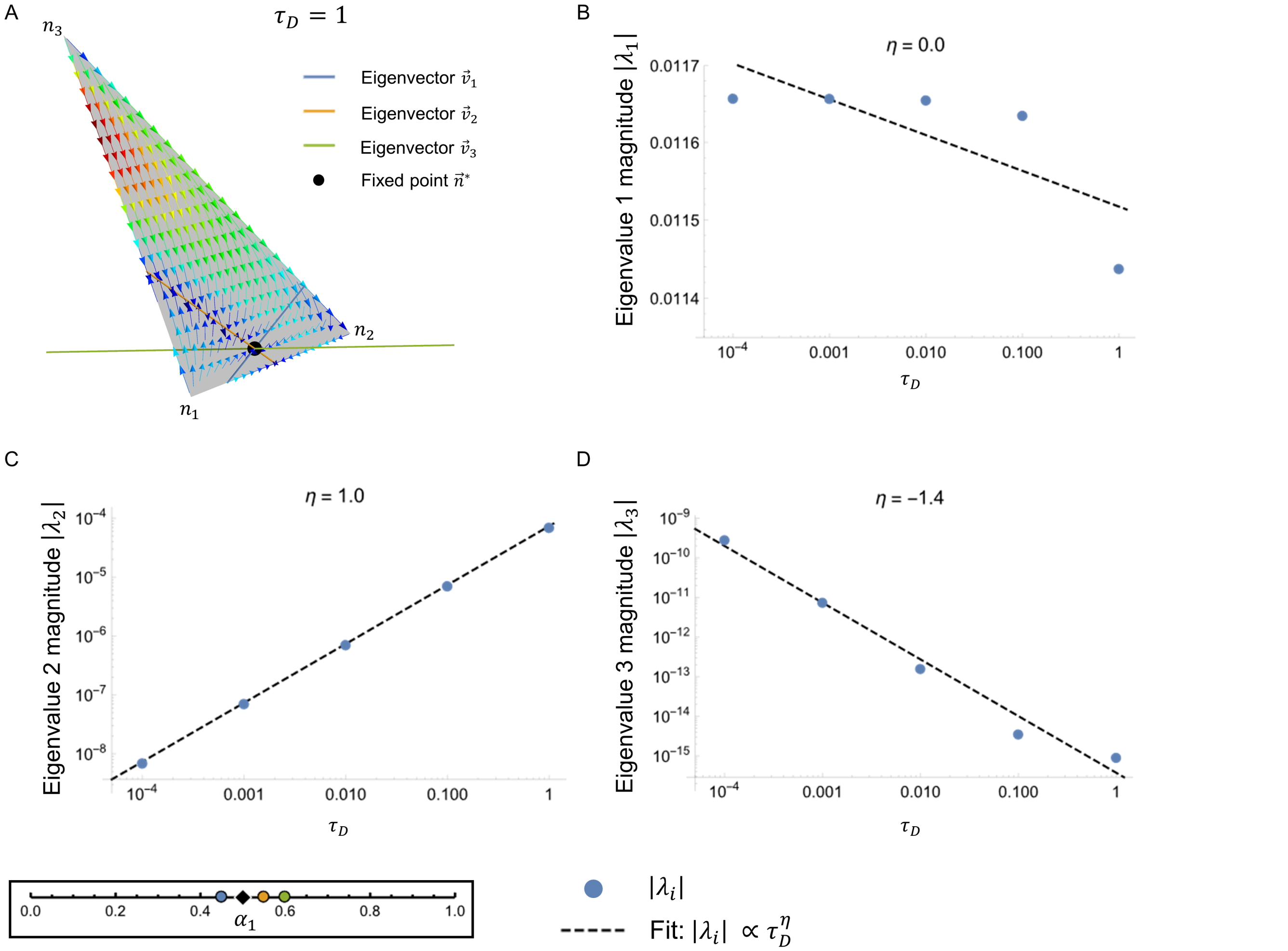


**Fig. S3.** Spatial territories lead to a slow manifold of dimension $m - p$ for dynamics in population space. (*A*) Trajectories in population space for a three-way competition. (Inset: strategies and supply). The direction and color of the arrows show the direction and magnitude, respectively, of $dn_\sigma/dt$. The eigenvector directions are obtained from numerical linear stability analysis around the fixed point. (*B*) The magnitude $|\lambda_1|$ of the first eigenvalue as $\tau_D$ is varied, with a fit (dashed line) to $|\lambda_1| \propto \tau_D^\eta$. This eigenvalue corresponds to $\vec{v}_1$ in *A*, and depends only weakly on $\tau_D$. (*C*) Same as *B* but for the eigenvalue of the slow manifold $\vec{v}_2$. Here the relaxation time $1/|\lambda_2| \propto 1/\tau_D$. (*D*) Same as *B* and *C*, but for $|\lambda_3|$. This eigenvalue describes stability to perturbations in the overall population size $L$.

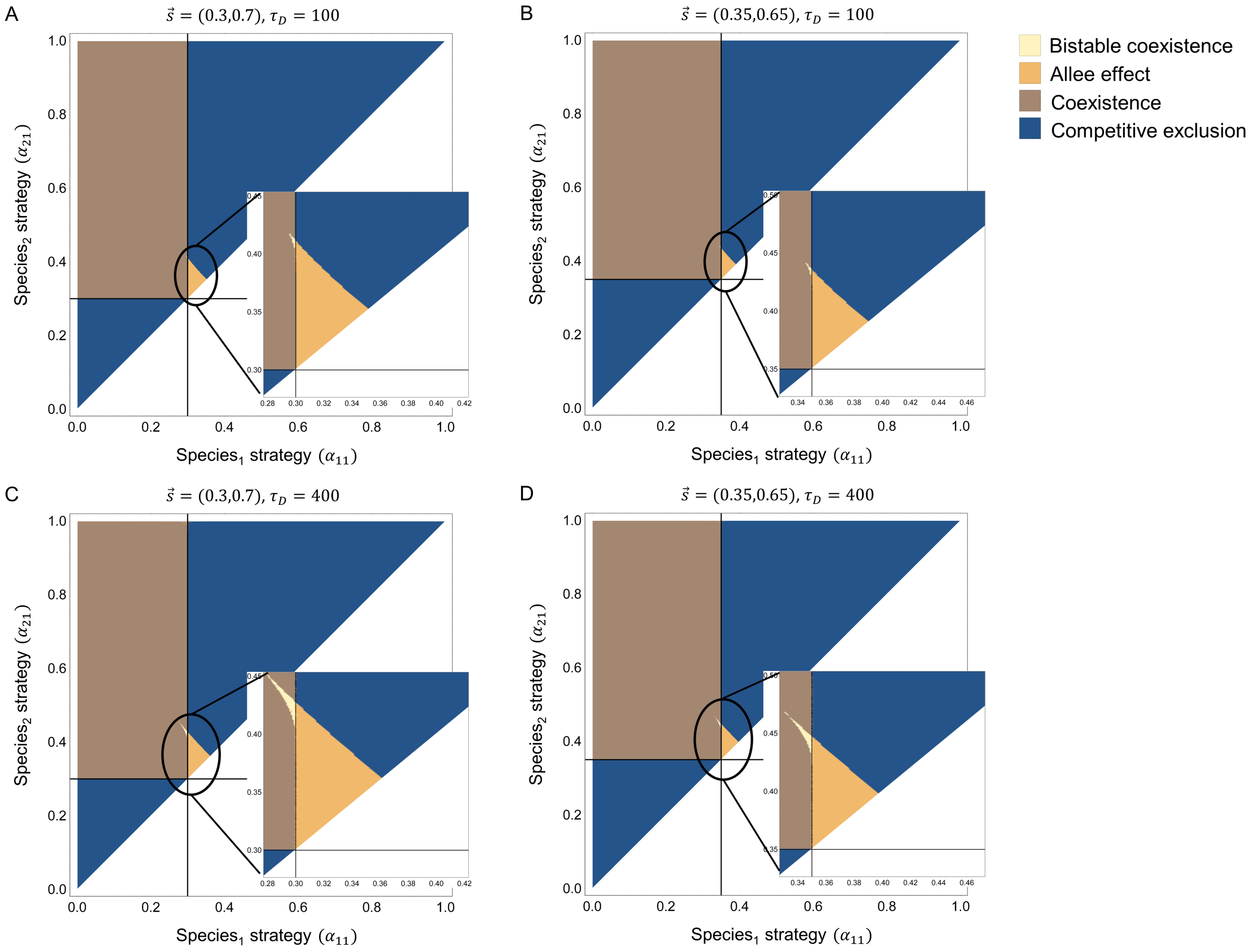


**Fig. S4.** Spatial structure leads to multistability and the Allee effect. (*A*) Classification of steady states depending on the strategies of two species competing for two resources, with $\vec{s} = (0.3, 0.7)$ and $\tau_D = 100$. (*B*) Same as *A*, but with $\vec{s} = (0.35, 0.65)$. The regions do not move relative to the point where both strategies match the supply, but they shrink. (*C*) Same as *A*, but for $\tau_D = 400$. The regions of strategies alternative stable states grow with increasing diffusion time. (*D*) Same as *B*, but for $\tau_D = 400$. Again, the strategy regions of alternative steady states grow.

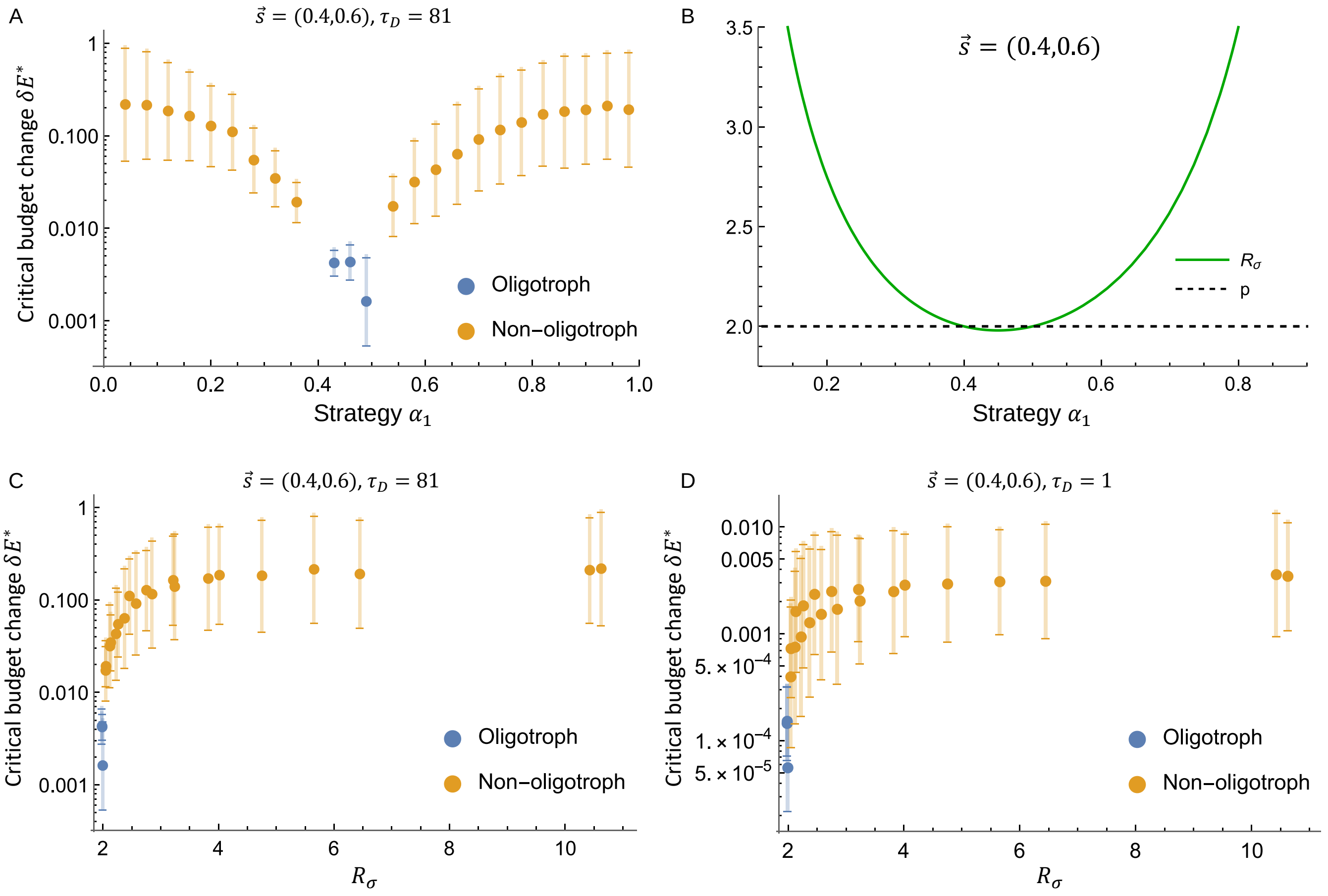


**Fig. S5.** In the spatial model, diversity beyond the competitive-exclusion limit persists for unequal enzyme budgets. This is because small changes in the total enzyme budget $E$ turn oligotrophs into non-oligotrophs, but the reverse requires large changes in $E$. (*A*) The change in total enzyme budget necessary to change final diversity depends on the strategy $\alpha_1$. For oligotrophs, $\delta E^*$ is the *decrease* in $E$ required for two more species to survive compared to the corresponding case with equal enzyme budgets. For non-oligotroph strategies, $\delta E^*$ is the *increase* in $E$ required for two fewer species to survive compared to the case of equal enzyme budgets. For each $\alpha_1$, $\delta E^*$ is averaged over 100 random sets of nine other species without oligotrophs $(\exp\left(\langle \log \delta E^* \rangle \pm SD\right)$, where $SD$ is the standard deviation of $\log \delta E^*$). (*B*) $R_\sigma = \sum_i \frac{S_i}{\alpha_{\sigma i}}$ as a function of strategy $\alpha_{\sigma i}$. Oligotroph strategies are very close to $R_\sigma = p$, while typical non-oligotroph strategies are far from $R_\sigma = p$. (*C*) Critical budget change as a function of $R_\sigma$. $\delta E^*$ increases rapidly outside the oligotroph region $R_\sigma < p$ before leveling off. (*D*) The same as *C*, but for smaller $\tau_D$. $\delta E^*$ has more variation, but the critical magnitude of $\delta E^*$ is still much smaller for oligotrophs than for non-oligotrophs.

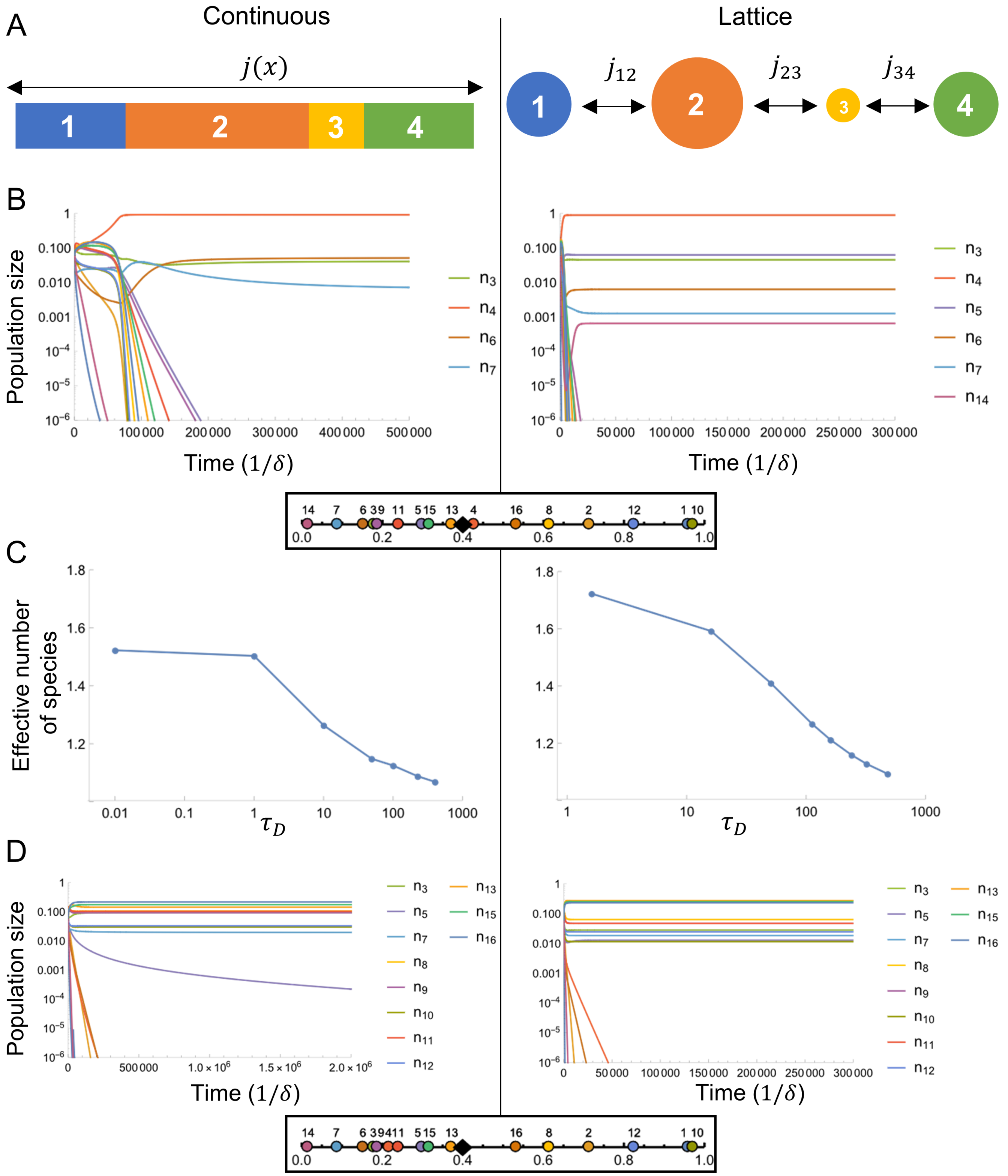


**Fig. S6.** A lattice territory model recapitulates the biodiversity patterns of the continuous territory model. (*A*) *Left:* In the continuous model of the main text, species occupy non-overlapping territories in a one-dimensional space of length $L$, and the nutrient flux $j(x)$ is continuous over $L$. *Right:* In the lattice model defined in Section 8, each territory occupies a lattice site. The nutrient environment in each territory is well-mixed, and territories on neighboring lattice sites $\sigma$ and $\sigma'$ exchange nutrients with flux $j_{\sigma\sigma'}$. (*B*) 16 species competing for 2 resources. In the well-mixed model, all 16 initial species would coexist, but in both the continuous and lattice models, most species go extinct. $n_T = 1$ in both models, corresponding to $\tau_D = 1$ for the continuous model and $\tau_D = 16$ for the lattice model (see Eq. 43). (Inset) Strategies $\vec{\alpha}_\sigma$ and resource supply $\vec{s} = (0.4, 0.6)$. (*C*) Effective number of species $M$ for various $\tau_D$. Same strategies and supply as *B*. (*D*) Same as *A*, but with sole oligotroph strategy $\vec{\alpha}_4$ changed to a non-oligotroph. (Inset) Strategies $\vec{\alpha}_\sigma$ and resource supply $\vec{s} = (0.4, 0.6)$.

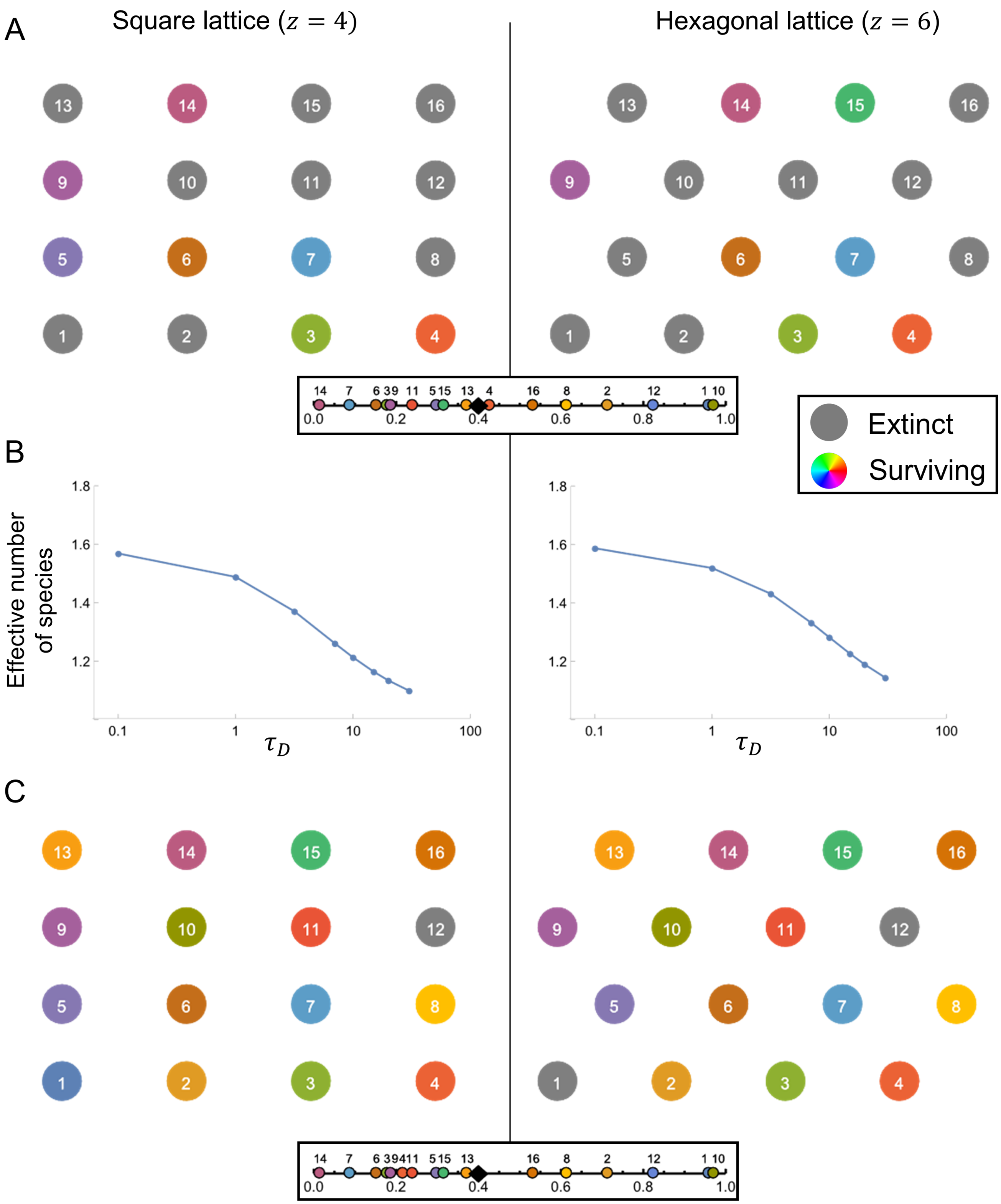


**Fig. S7.** The lattice territory model in two spatial dimensions displays biodiversity patterns similar to the one-dimensional continuous model. (*A*) Coexistence of species at steady state on a square lattice with four neighbors (*Left*) or on a hexagonal lattice with six neighbors (*Right*), for $\tau_D = 1$. Colored and gray circles denote surviving and extinct populations, respectively. A population is considered extinct if $n_\sigma/n_T < 10^{-7}$. (Inset) Strategies $\vec{\alpha}_\sigma$ and resource supply $\vec{s} = (0.4, 0.6)$ (same as in Fig. S6*B* and *C*). (*B*) Effective number of species $M$ on the square lattice (*Left*) and hexagonal lattice (*Right*), as a function of $\tau_D$. Same strategies and supply as *A*. (*C*) Same as *A*, but with sole oligotroph strategy $\vec{\alpha}_4$ changed to a non-oligotroph. (Inset) Strategies $\vec{\alpha}_\sigma$ and resource supply $\vec{s} = (0.4, 0.6)$ (same as in Fig. S6*D*).

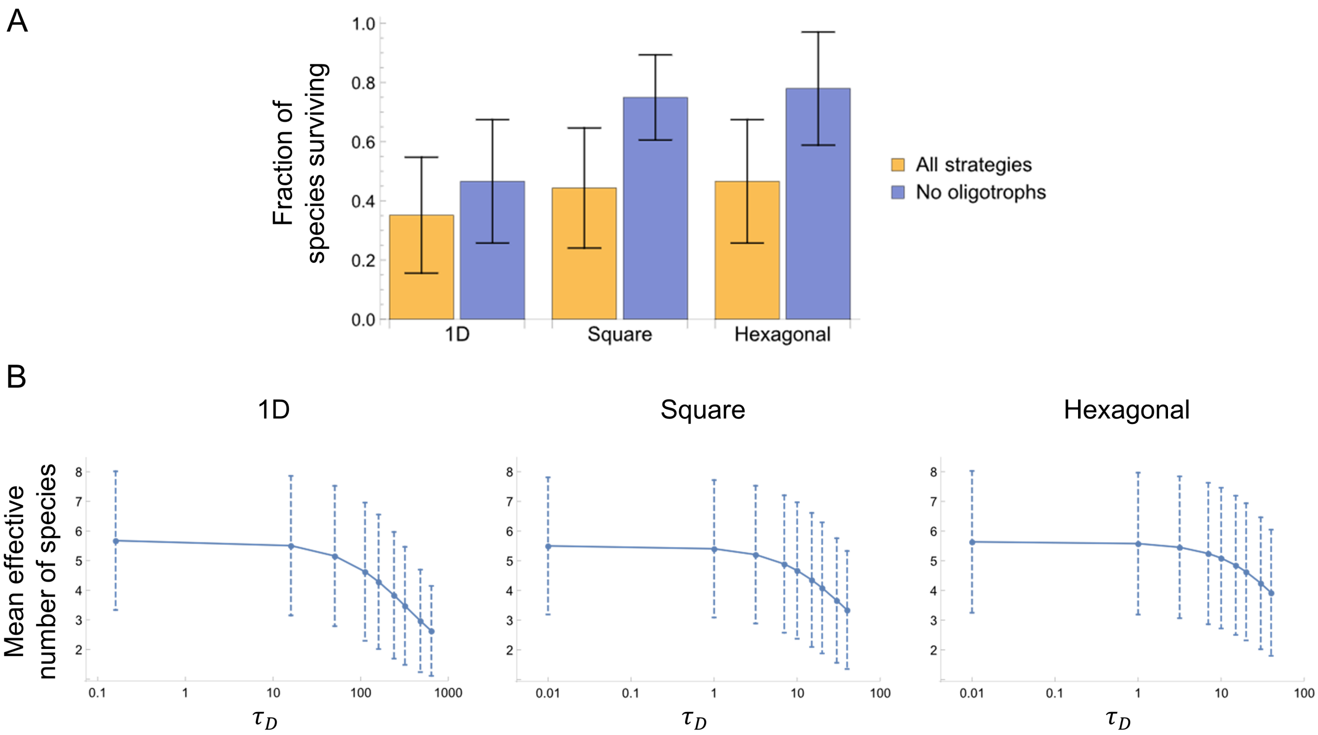


**Fig. S8.** The biodiversity patterns exhibited by the lattice territory model in Fig. S6 and S7 are generic for random sets of strategies. (*A*) Fraction of species surviving at steady state for three lattice geometries. A population is considered extinct if $n_\sigma/n_T < 10^{-6}$ (mean $\pm$ SD for 300 sets of 16 strategies with $\vec{s} = (0.4, 0.6)$). We use $n_T = 1$ in each geometry, yielding $\tau_D = 16$ for the 1D lattice and $\tau_D = 1$ for the square and hexagonal lattices. (*B*) Effective number of species at steady state for three lattice geometries, with oligotroph strategies permitted (mean $\pm$ SD for 300 sets of 16 strategies at each $\tau_D$, with $\vec{s} = (0.4, 0.6)$).

### References

1. Posfai A, Taillefumier T, Wingreen NS (2017) Metabolic trade-offs promote diversity in a model ecosystem. *Physical Review Letters* 118(028103).
